# Digital twins for understanding mechanisms of learning disabilities: Personalized deep neural networks reveal impact of neuronal hyperexcitability

**DOI:** 10.1101/2024.04.29.591409

**Authors:** Anthony Strock, Percy K. Mistry, Vinod Menon

## Abstract

Learning disabilities affect a significant proportion of children worldwide, with far-reaching consequences for their academic, professional, and personal lives. Here we develop digital twins – biologically plausible personalized Deep Neural Networks (pDNNs) – to investigate the neurophysiological mechanisms underlying learning disabilities in children. Our pDNN reproduces behavioral and neural activity patterns observed in affected children, including lower performance accuracy, slower learning rates, neural hyper-excitability, and reduced neural differentiation of numerical problems. Crucially, pDNN models reveal aberrancies in the geometry of manifold structure, providing a comprehensive view of how neural excitability influences both learning performance and the internal structure of neural representations. Our findings not only advance knowledge of the neurophysiological underpinnings of learning differences but also open avenues for targeted, personalized strategies designed to bridge cognitive gaps in affected children. This work reveals the power of digital twins integrating AI and neuroscience to uncover mechanisms underlying neurodevelopmental disorders.

---

The early years of childhood are pivotal for the development of foundational academic and cognitive skills, a process marked by significant individual variability among children ^1-3^. Among the essential skills, mathematical proficiency stands out as a particularly challenging area for a subset of children ^4,5^. Mathematical Learning Disabilities (MLD), which affects about 5 to 20 percent of the global population, manifests as diminished problem-solving abilities when benchmarked against peers with similar age and intelligence ^5-7^. The repercussions of MLD extend beyond academic challenges, impacting long-term socioeconomic status, including employment prospects and health outcomes ^8-10^. Despite extensive research over the past two decades, the neurobiological underpinnings of MLD remain elusive. Harnessing artificial intelligence (AI) models that capture individual variability and can serve as digital twins^11-13^ – capturing critical components of a biophysical system while allowing for in silico experimentation – offers new promise. Here we introduce a novel approach by developing a biologically plausible ^14,15^ personalized deep neural network (pDNN). These personalized networks are tailored to mirror individual behavioral patterns and evaluated against neuroimaging data obtained from the same individuals, thus serving as digital twins on which further experimentation and analysis of neuronal mechanisms can be performed, which would be difficult to achieve non-invasively in children. Our goal is to uncover the hidden neural mechanisms and representations that underpin individual differences in mathematical cognition, offering new insights into the complex interplay between brain function and MLD.

MLD is marked by significant challenges in arithmetic problem-solving, a foundation for developing advanced mathematical concepts ^16,17^. Children with MLD often struggle with basic arithmetic operations, such as addition and subtraction ^5,18^. These difficulties are not just limited to slower processing speeds but encompass lower accuracy and use of less efficient problem-solving strategies ^19-21^. Although various cognitive factors have been implicated ^5^, a unifying, biologically plausible model for MLD has, however, been elusive. Neuroimaging has played a crucial role in uncovering the brain basis of MLD ^6,17,22-26^, with significant dysfunctions identified in regions critical for numerical cognition, such as the intraparietal sulcus (IPS) ^27,28^. Moreover, abnormalities extend to a broader network involved in visual and visuospatial processing, suggesting MLD as a multifaceted neural dysfunction ^23,24,29-32^. However, our understanding of the underlying neural mechanisms, which are vital for overcoming the challenges posed by MLD, remains limited.

Recent results have identified reduced behavioral and neural differentiation between distinct numerical operations in children with MLD ^21^. This suggests less efficient neural processing, characterized by over engagement of brain circuits beyond levels typically needed for task performance. Research has also shown that such impairments often extend into adulthood in individuals with dyscalculia, highlighting the persistent nature and long-term effects of MLD^33^.

Aligned with inferences about less efficient neural processing, recent studies have identified hyperactive brain patterns and hyperconnectivity ^18,34^ in key cognitive regions among children with MLD, suggesting an over-synchronization of neural networks essential for numerical cognition. Examination of the amplitude of intrinsic low-frequency fluctuations, a proxy measure for regional neural activity, has extended our understanding of dysfunctional neural circuits associated with poor math abilities ^34^. It has been discovered that children with MLD exhibit greater signal fluctuation across multiple brain regions, a finding indicative of neural hyper-excitability ^34^. This result has been further substantiated by reports that parietal and hippocampal hyperconnectivity is associated with low math achievement in adolescence ^35^. Additionally, hyperactivity is associated with greater intrinsic functional connectivity between multiple cortical regions. Aligned with this pattern of hyperactivity, magnetic resonance spectroscopy investigations have pointed to differential Glutamate and GABA concentration, indicative of excitation/inhibition (E/I) imbalances, in children with poor math abilities ^36,37^, in expert math calculation (reduced frontal E/I balance) ^38^. GABA and glutamate in the IPS have also been shown to explain individual variability in mathematical achievement levels^39^, and in test anxiety levels in early childhood ^40^. Individuals lacking in mathematical education have also been shown to demonstrate lower inhibition in brain regions relevant to reasoning and learning^36^. Recent studies which attempt to rectify E/I imbalance by using neurostimulation have shown that E/I balance modulates the amount of benefit that can be obtained from neurostimulation ^41-43^, and can be a marker for neurostimulation based efficacy and learning^41^. Despite these advances, the neurophysiological mechanisms underlying MLD, the sufficiency of establishing E/I imbalances as contributing factors towards MLD, and the mechanisms via which such imbalances may cause learning difficulties, remain speculative, primarily due to the correlative nature of brain imaging studies.

Digital twins, operationalized here via pDNN models, provide a novel lens for addressing crucial knowledge gaps in the neurophysiology of mathematical cognition and learning disabilities. DNNs have demonstrated significant success in modeling a variety of cognitive functions including number sense ^14,44,45^, word reading ^46^, object recognition ^47,48^, and sentence processing ^49^, yet their application in understanding learning disabilities has been scant, primarily due to lack of theoretically motivated approaches for introducing individual differences in DNN behavior. Motivated by the potential role of E/I imbalances in contributing towards learning differences, we focus on using neural excitability, as a key theoretical mechanism by which individual differences can be introduced into DNN models.

This study thus introduces a novel personalized pDNN framework (**Figure 1A**) aimed at modelling individual behavioral performance and linking it to functional brain imaging data, to elucidate the specific neurobiological dysfunctions associated with MLD. Specifically, we aimed to model behavioral and neural deficits in numerical problem-solving using addition and subtraction, two fundamental operations crucial for early numerical problem-solving proficiency ^5,23,50^. Behavioral studies suggest that performance on tasks problems involving addition and subtraction operations is characterized by significant individual differences in problem-solving abilities in children, and that children with MLD are impaired on both ^5,18,50,51^. By leveraging biologically plausible DNNs ^14^, and integrating behavioral with neuroimaging data, we explored individual variability and the neural correlates of arithmetic learning, thereby contributing to a deeper understanding of the neural underpinnings of core numerical problem-solving skills essential for early cognitive development.

**Figure 1.**
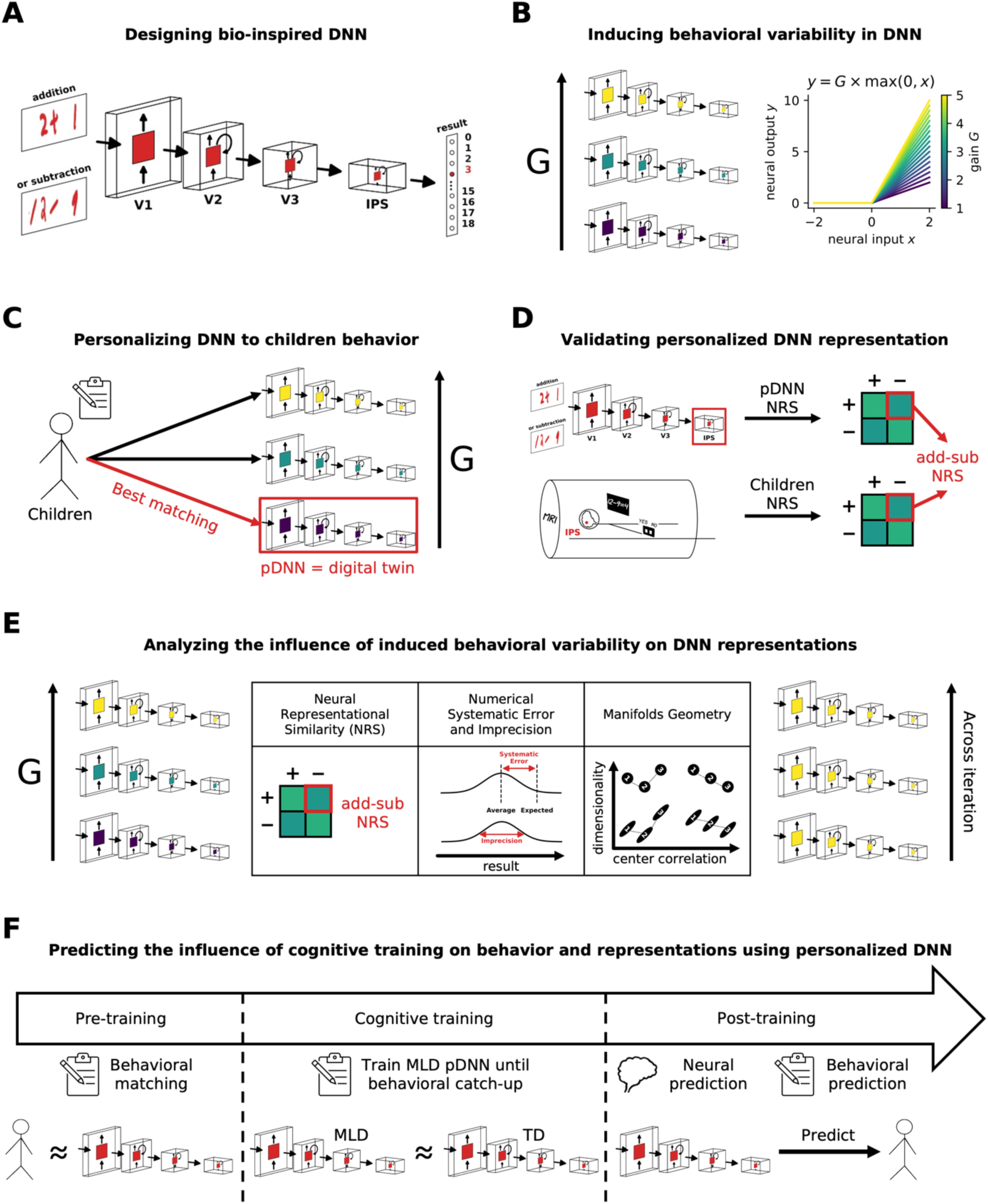
Design and analysis of personalized deep neural networks (pDNNs) for modeling numerical cognition and learning disabilities. **A.** Schematic depicting a biologically inspired deep neural network (DNN) model mimicking the dorsal visual pathway involved in numerical cognition. The model is trained to solve visually presented addition and subtraction problems. **B.** Schematic illustrating the modulation of neural excitability in the model, based on brain imaging studies suggesting a link between excitation/inhibition (E/I) balance and learning capabilities. Neural excitability is measured as the gain *G*. **C.** Creating digital twins – personalized DNNs (pDNNs) that match individual children’s performance levels (assessed by *NumOps*) – by adapting neural excitability. Based on previous brain imaging studies, we hypothesize that neural hyper-excitability (i.e., higher *G*) is a plausible mechanism underlying mathematical learning disabilities (MLD) compared to typically developing (TD) children. **D.** Validation of pDNNs by verifying whether they exhibit the same representational deficits observed in brain imaging studies, namely lower neural differentiation of numerical problems measured with Neural Representational Similarity (NRS). **E.** Schematic depicting further analyses of how multiple behavioral and representational aspects of the model evolve during training and with varying levels of neural excitability. **F.** Investigation of the influence of additional training on pDNNs in the MLD group and the associated changes in latent neural representations. This analysis aims to uncover the potential for remediation and the neural mechanisms underlying the improvements in numerical cognition following targeted training.

Our study had five primary objectives. Our first objective was to investigate E/I imbalance, characterized as neural excitability, as a potential and sufficient neurobiological mechanism (**Figure 1B**) contributing to cognitive performance deficits in children with MLD. Neural excitability and E/I balance is a fundamental aspect of neural processing, playing a crucial role in shaping neural network dynamics, learning, and cognitive function ^52-55^. Moreover, as reviewed above, task-related hyperactivation and intrinsic hyper-fluctuations observed in children with MLD suggest that E/I imbalance may be a key underlying neurophysiological mechanism. The concept of E/I imbalance as a putative neural mechanism underlying learning disabilities is suggested by both theoretical models and empirical research ^41,43,53,55-57^. E/I balance is a fundamental aspect of neural processing, playing a crucial role in shaping neural network dynamics and cognitive function ^52-55^. We therefore probed whether neural excitability mechanisms can induce systemic individual differences in the arithmetic problem-solving behaviors of biologically inspired DNNs. We hypothesized that varying levels of neural excitability could capture relatively monotonic changes in learning, behavioral patterns, and latent neural representations, and show meaningful structure in terms of how neural representations evolved over the network hierarchy with changing excitability levels.

Our second objective was to construct pDNNs, where we could tune the neural excitability levels to match the individualized learning profiles of children performing a similar arithmetic task (**Figure 1C**) and capture the influence of such varying neural excitability on individual variations in behavioral task performance. We demonstrate that by manipulating neural excitability parameters we can, for the first time, match personalized networks to best represent the behavioral aspects of individual children, thus creating digital twins for both typically developing children (TD) and children with MLD. We were thus able to obtain a set of TD pDNNs and MLD pDNNs and evaluate differences between these sets of networks.

Our third objective was to use fMRI data from children to assess whether pDNNs that were behaviorally matched to individual children’s profiles were predictive of individual differences in neural representations observed in empirical data (**Figure 1D**). We hypothesized that behaviorally matched pDNNs would reasonably predict aberrant neural representations in children with MLD. If successful, this would validate the concept of *in silico* digital twins, and insights obtained from pDNNs could be used not just to draw inferences about how neural excitability affects different aspects of neural representations, but also make individualized predictions.

Our fourth objective was to analyze the neural data from the best matched pDNNs to extract insights about how latent neural representations varied with changing neural excitability (**Figure 1E**). Our hypothesis was that pDNNs matched to MLD children would show differences in such latent neural representations compared to pDNNs matched to TD children, thus developing the link between differences in E/I imbalances and latent neural representations of mathematical problem solving. This would reveal aspects of information processing deficits, how these are distributed across the hierarchical structure representing the dorsal visual stream and IPS, and how the representational geometry of distributed neural measures such as neural manifold capacity, structure, and dimensionality, impact learning deficits^58,59^.

Our fifth objective was to explore whether additional training can normalize behavioral performance and neural representation patterns in children with MLD to levels observed in TD children (**Figure 1F**). We hypothesized that E/I imbalances characterized by neural excitability would slow down but not put a hard constraint on learning. We determined how much additional training would be required for MLD pDNNs to match TD pDNNs levels of performance, and whether such training was accompanied by changes in latent neural representations like those seen in TD pDNNs. This approach aimed to provide insights into the adaptive capacity of the neural processes in MLD, potentially informing future intervention strategies to address disabilities linked to E/I imbalances.

Our findings demonstrate the significant potential of digital twins, operationalized via pDNNs, in uncovering latent neurobiological mechanisms underlying individual differences in children’s behavior and learning. We show that pDNNs can simulate and assess the impact of neural excitability on cognitive performance, creating a bridge between AI, computational neuroscience, and empirical brain imaging studies. Our approach provides a novel framework for linking neural network models with cognitive neuroscience studies in human participants.

## Results

Figure 1 shows the study design, data used and critical steps of our analysis strategy. We first developed a pDNN model for numerical problem-solving tasks involving addition and subtraction operations. Our pDNN models were constructed using a biologically inspired model of the dorsal visual pathway based on the network architecture and physiological parameters of CORnet-S ^14,15^. This neural architecture, comprising cortical layers V1, V2, V3, and intraparietal sulcus (IPS), has been shown to characterize how neural representations can change with numerosity training, and how learning can reorganize neuronal tuning properties at both the single unit and population levels^14^. Importantly, such models have been able to capture learning driven changes from logarithmic to cyclic and linear mental number lines that are characteristic of number sense development in humans^14^. The models were utilized in their raw form, without any pre-training (Figure 1A). The training problems were similar to those used in our fMRI study with children ^21^, incorporating images of handwritten arithmetic operations with results spanning natural numbers from 0 to 18, designed to mimic the diversity of handwriting children might encounter in educational settings (see **Methods** for details). This approach ensured the robustness and generalizability of the pDNNs in real-world learning scenarios.

Our empirical data pool consisted of 45 children, aged 7 to 9, from second and third grades, who performed numerical problem-solving tasks analogous to those processed by the pDNNs. Out of the 45 participants, 21 children were identified with MLD based on their NumOps scores on standardized WIAT-II ^60^ test subscores. The remaining 24 children were considered TD and served as the control group. The two groups did not differ on age, full-scale IQ, or reading abilities (**Table SI 1**). All children solved addition and subtraction problems during fMRI scanning. In the addition task, they were presented with an equation (e.g., “3 + 4= 8”), and were asked to indicate, via a button press, whether the presented answer was correct. 36 addition problems were presented, with 50% paired with correct answers and 50% with incorrect answers. A similar procedure was used for subtraction problems. Further details of the study protocol and design are presented in the **Methods** section and in previous studies ^21^.

### Tuning neural excitability produces individual differences within pDNNs

The basic pDNN models were personalized and individual differences in task performance were simulated by tuning the neural excitability (neural gain parameter *G*, Figure 1B). Specifically, we varied this parameter from 1.0 to 5.0 in steps of 0.25. Each of these 17 models with different levels of neural gain was trained on the same set of problems for a fixed number of iterations. At the outset, all pDNN models operated at chance levels of accuracy (approximately 5%, with possible answers ranging from 0 to 18), ensuring a uniform starting point. We hypothesized that heightened neural excitability could potentially impede the learning process, either by decelerating the rate of learning or by limiting the ultimate proficiency attainable by the pDNN. In either case, under this hypothesis, any fixed number of iterations within a certain bound would result in a situation where pDNNs with lower neural excitability would have a higher level of behavioral accuracy, allowing us to match different neural excitability levels to children with different mathematical achievement levels.

To assess the role of neural excitability on the learning efficiency of our pDNNs, we focused on the number of training iterations required for the models to achieve a 95% accuracy threshold. This threshold was indicative of mastery in solving the addition and subtraction problems used in our study. We found that models with the lowest level of neural excitability were able to achieve this threshold in about 1400 iterations, but after about 3800 training iterations, all models achieved an accuracy exceeding 95%, suggesting that pDNNs models with a wide range of neural excitability were able to learn to solve addition and subtraction problems with high reliability (Figure 2A). These results underscore the potential of pDNNs for simulating numerical problem-solving and learning in children (Figure 2A).

**Figure 2.**
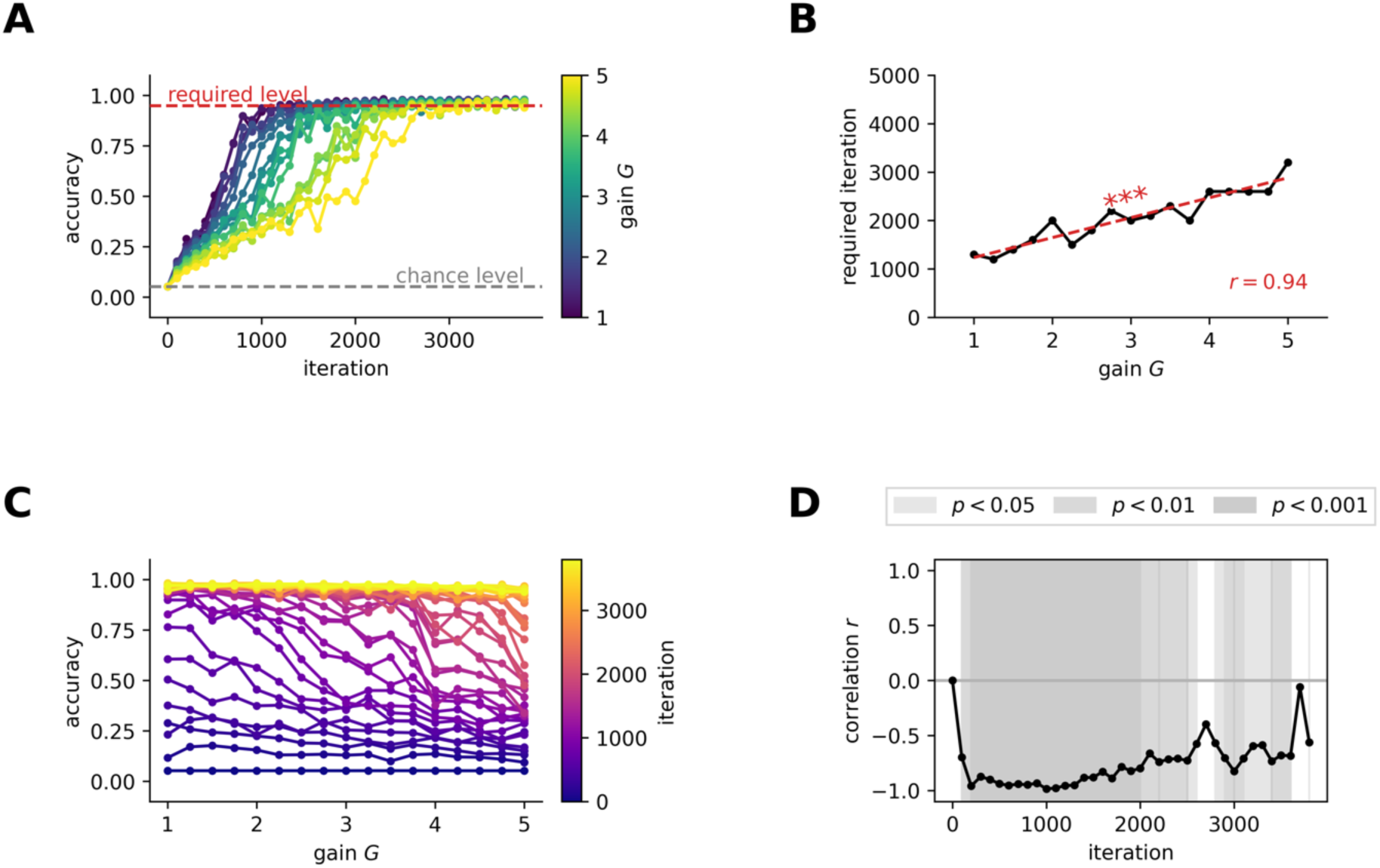
Neural hyper-excitability reduces learning speed and accuracy in DNNs. **A.** Learning trajectories of DNN models with different levels of neural excitability, measured as neural gain *G*. Neural gain *G* is represented by color, varying from blue (*G* = 1) to yellow (*G* = 5). As neural excitability increases, the progression in accuracy across learning iterations slows down, indicating a slower learning rate. **B.** Number of iterations required to reach an accuracy of 95% for different values of neural gain *G*. The number of iterations needed to reach the 95% accuracy benchmark consistently increases with neural excitability (*r* = 0.94, *p* < 1e-7), demonstrating that heightened excitability impairs learning efficiency. **C.** Changes in DNN test accuracy with neural gain *G* for different iterations, represented by color ranging from blue (iteration 0) to yellow (iteration 3800). Higher neural gain values are associated with lower test accuracies across all iterations, suggesting that hyper-excitability hinders the model’s ability to generalize to new problem sets. **D.** Correlation between DNN test accuracy and neural gain *G* across iterations. The negative correlation between test accuracy and neural gain remains consistent throughout the training process. This indicates that the detrimental effect of hyper-excitability on learning and generalization persists across the training trajectory. *** *p* < 0.001.

However, our findings revealed distinct learning trajectories across pDNNs with varying levels of neural excitability. As excitability increased, we observed a slower progression in accuracy across learning iterations. The number of iterations required to reach the 95% accuracy benchmark increased consistently with neural excitability (*r* = 0.94, *p* < 10^−7^, Figure 2B).

After 200 iterations, the relationship between neural excitability and model performance revealed a strong inverse correlation ( *r* = −0.96, *p* < 10^−8^, Figures 2C-D). This negative correlation remained consistent throughout the training process and only began to diminish when pDNNs approached peak performance levels (Figures 2C-D).

These results demonstrate that increased neural excitability in pDNNs does not impede the models’ ability to eventually achieve high levels of accuracy in arithmetic tasks, but does lead to a slower learning rate, and that tuning the neural excitability of pDNNs can thus produce individual differences in behavioral performance at any fixed learning iteration (analogous to a certain level of training in humans).

### pDNNs tuned based on neural excitability can be behaviorally matched to individual children – MLD matched pDNNs demonstrate higher neural excitability

Next, we fine-tuned pDNNs to the specific mathematical achievement level of each child participant. By adjusting the neural excitability values in the pDNNs, we aimed to closely approximate the mathematical problem-solving behaviors exhibited by children. Matching was carried out using ‘behavioral distance’ (see **Methods** for details), a measure of the degree of alignment between normalized values of each child’s mathematical achievement scores and the pDNN performance. Since the pDNNs ‘experienced’ a much wider range of arithmetic problems compared to children in the specific task, the matching was done by comparing normalized accuracy of the pDNNs across all addition and subtraction problems, with the normalized *NumOps* ^60^ score of children. This ensured that the comparison was robust and ecologically valid, and that since the comparison was not based on direct task-proximal measures or specific problem subsets, the neural predictions would be generalizable. By evaluating every 100^th^ learning iteration of the pDNNs, we determined the best-fitting neural excitability (gain value *G*) for each individual child at each iteration. We then identified the specific learning iteration at which the best-fitting pDNNs for each child most accurately reflected the behavioral achievement levels (**Methods**) of the children (Figure 3A) on an average. This analysis identified iteration 800 as optimal for capturing the full spectrum of individual differences in the behavioral performance of mathematical problem-solving seen in child participants.

**Figure 3.**
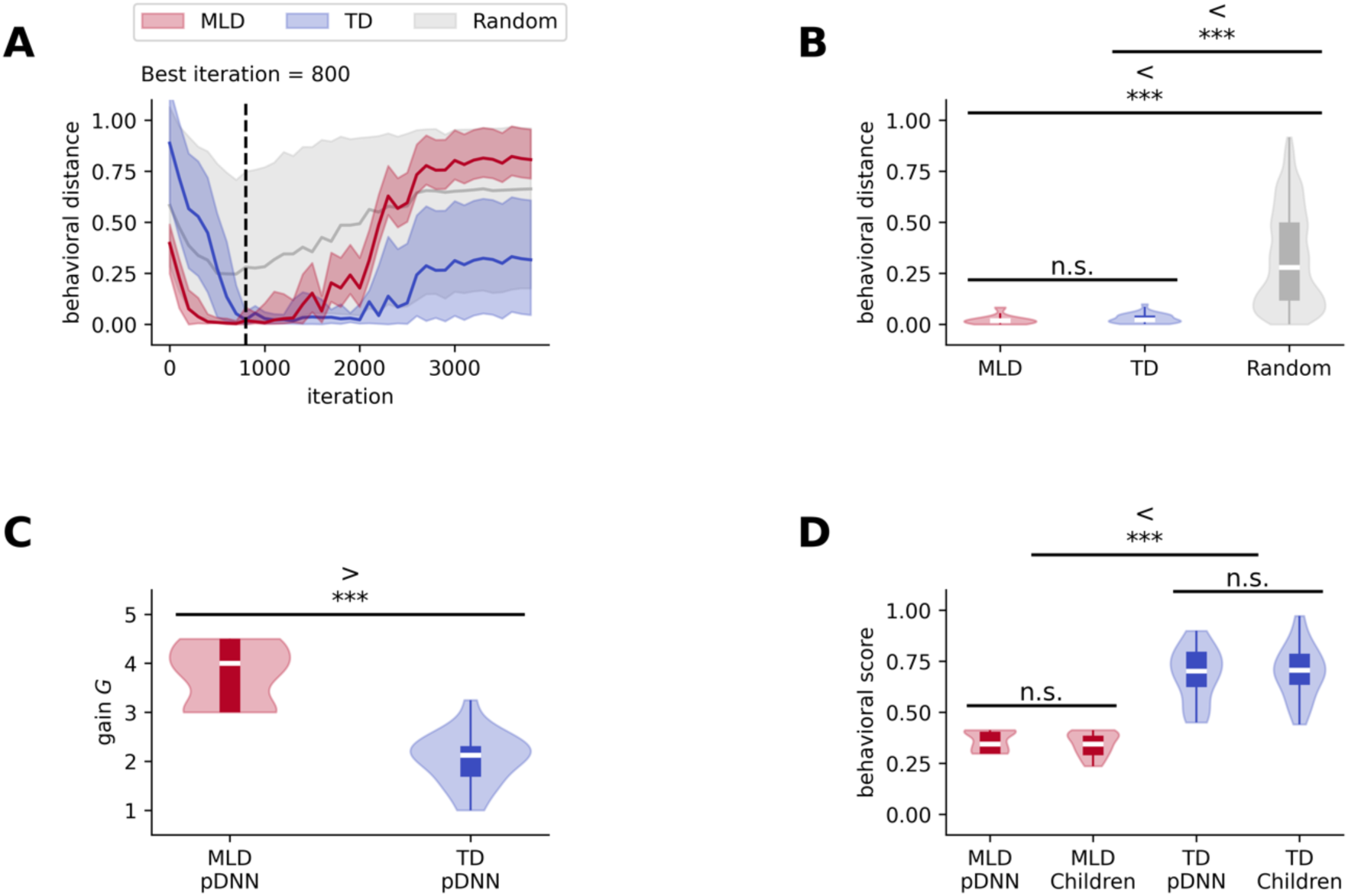
Personalized deep neural networks (pDNNs) tuned for neural excitability capture individual differences in children’s math performance and serve as *in silico* digital twins. **A.** A distance metric between pDNN model accuracy and children’s normalized math scores across training iterations was used to identify the best matching point (iteration 800, dotted black line). The red and blue lines represent the average distance for pDNNs tuned to match children with mathematical learning disabilities (MLD) and typically developing (TD) children, respectively. Gray lines show the average distance for pDNNs with randomly assigned neural excitability levels, serving as a control. Shaded areas denote the range between the 5th and 95th percentiles across children in each group. **B.** Distribution of the distance metric at iteration 800 for MLD (red) and TD (blue) groups, compared to the random control (gray). The distance metric for both MLD and TD groups is significantly lower than the controls (MLD: *p < 1e-38*; TD: *p < 1e-46*), indicating a strong match between pDNNs and children’s behavioral performance. **C.** Distribution of the best-matched neural excitability levels (gain *G*) at iteration 800 for MLD and TD groups. The neural excitability levels are significantly higher for pDNNs matched to MLD children compared to those matched to TD children, suggesting that higher neural excitability is associated with math learning difficulties. **D.** Comparison of behavioral performance distributions between pDNNs and children at iteration 800. The normalized behavioral scores of MLD children and their matched pDNNs do not differ significantly, and the same holds for TD children and their matched pDNNs. Both MLD children and their matched pDNNs show significantly lower behavioral scores compared to TD children and their matched pDNNs, respectively (children: *p < 1e-11*; pDNNs: *p < 1e-10*). These results demonstrate that pDNNs tuned for neural excitability accurately capture the individual differences in math performance observed in children. *** *p* < 0.001.

Figure 3A shows the 95% CI for behavioral distances between the children and their best matched pDNNs for the TD and MLD groups across different iterations. Iteration 800 was a good fit for both the TD and MLD groups, and the difference in behavioral distance (measure of fit) between groups was not significant (*t* = −1.09, *p* = 0.28, Figure 3B). We conducted a control analysis in which we randomly permuted the children’s behavioral scores across different neural gains at iteration 800. The resulting behavioral distance for these random permutations (*M* = 0.32, *SD* = 0.24) was significantly higher than pDNN fitted data at this iteration for both MLD (*M* = 0.02, *SD* = 0.02) (*t* = 48.07, *p* < 10^−38^, Figure 3B) and TD groups (*M* = 0.03, *SD* = 0.02) (*t* = 48.96, *p* < 10^−46^, Figure 3B). This result supports the finding that our model is well-calibrated to empirical data at iteration 800, thereby establishing it as the focal point for in-depth analysis in subsequent sections. Going forward, the best fitting excitability models at iterations 800 for TD and MLD groups respectively are collectively termed *TD pDNNs* and *MLD pDNNs* respectively and represent the digital twins for these groups. Figure 3C shows that, as we hypothesized, the gain of MLD pDNNs (*M* = 3.77, *SD* = 0.60) is higher than the gain of TD pDNNs (*M* = 2.05, *SD* = 0.52) (*t* = 9.96, *p* < 10^−11^, Figure 3C).

These findings demonstrate that pDNNs can be individually tailored to represent children’s varying levels of performance on mental arithmetic tasks, that neural excitability is a sufficient and key factor in this personalization of pDNNs, and that MLD pDNNs are associated with higher neural excitability than TD pDNNs. They also demonstrate that the personalized pDNNs have similar degrees of fit for both TD and MLD children, showing that the personalized models are not inherently biased and can effectively cover the full spectrum of individual differences.

### Behavioral patterns of pDNN models are good representations of empirical behavior of TD and MLD children in arithmetic tasks

Although pDNNs were individually matched by selecting the iteration that minimized average differences in normalized behavioral scores for each child, there was no guarantee that these behavioral scores would show a good absolute fit at this iteration. We evaluated whether pDNN performance accurately reflected the behavioral achievement patterns observed in MLD and TD children. We compared normalized performance scores of these two groups of children (*NumOps*) and the normalized accuracy of their corresponding pDNNs. Our statistical comparison revealed no evidence of significant differences between the normalized behavioral scores of MLD children (*M* = 0.35, *SD* = 0.06) and their matched MLD pDNNs (*M* = 0.36, *SD* = 0.05) (*t* = 0.98, *p* = 0.33, Figure 3D) and no evidence of significant differences between the scores of TD children (*M* = 0.67, *SD* = 0.13) and their matched TD pDNNs (*M* = 0.69, *SD* = 0.13) (*t* = −0.26, *p* = 0.80, Figure 3D).

Figure 3D shows that the behavioral scores of MLD pDNNs were significantly lower than TD pDNNs (*t* = −11.06, *p* < 10^−11^, Figure 3D), and are well aligned with our empirical findings of lower performance in children with MLD (*t* = −11.86, *p* < 10^−12^, Figure 3D).

These results further validate that the behavioral scores of pDNNs, when matched to individual math achievement levels, accurately reflect empirical data, affirming the utility of pDNNs in modeling the behavioral nuances of children with MLD and TD in mental arithmetic tasks.

### Neural representational similarity (NRS) between problem types increases with neural excitability in pDNN models

Our next goal was to investigate the effect of neural excitability on the pDNN NRS between addition and subtraction, two distinct numerical operations. This analysis was motivated by our empirical evidence of higher NRS between addition and subtraction problems in children with MLD, compared to TD children ^21^. This profile of less differentiated neural representations was particularly prominent in the IPS region in children.

We averaged the NRS between each individual pair of operations in the pDNN model and examined how this similarity changed with neural excitability *G* (Figures 4A-B show these for low and high neural excitability of *G* = 2.25 and *G* = 4.0). NRS was computed for each iteration, for each level of neural excitability, and within each layer of the pDNN (V1, V2, V3, IPS), as described in **Methods**. These results show that while the NRS between operations is similar for high and low gains in the lower V1 layer, the difference amplifies over the layer hierarchy and shows significant differences between high and low gains in the IPS layer.

**Figure 4.**
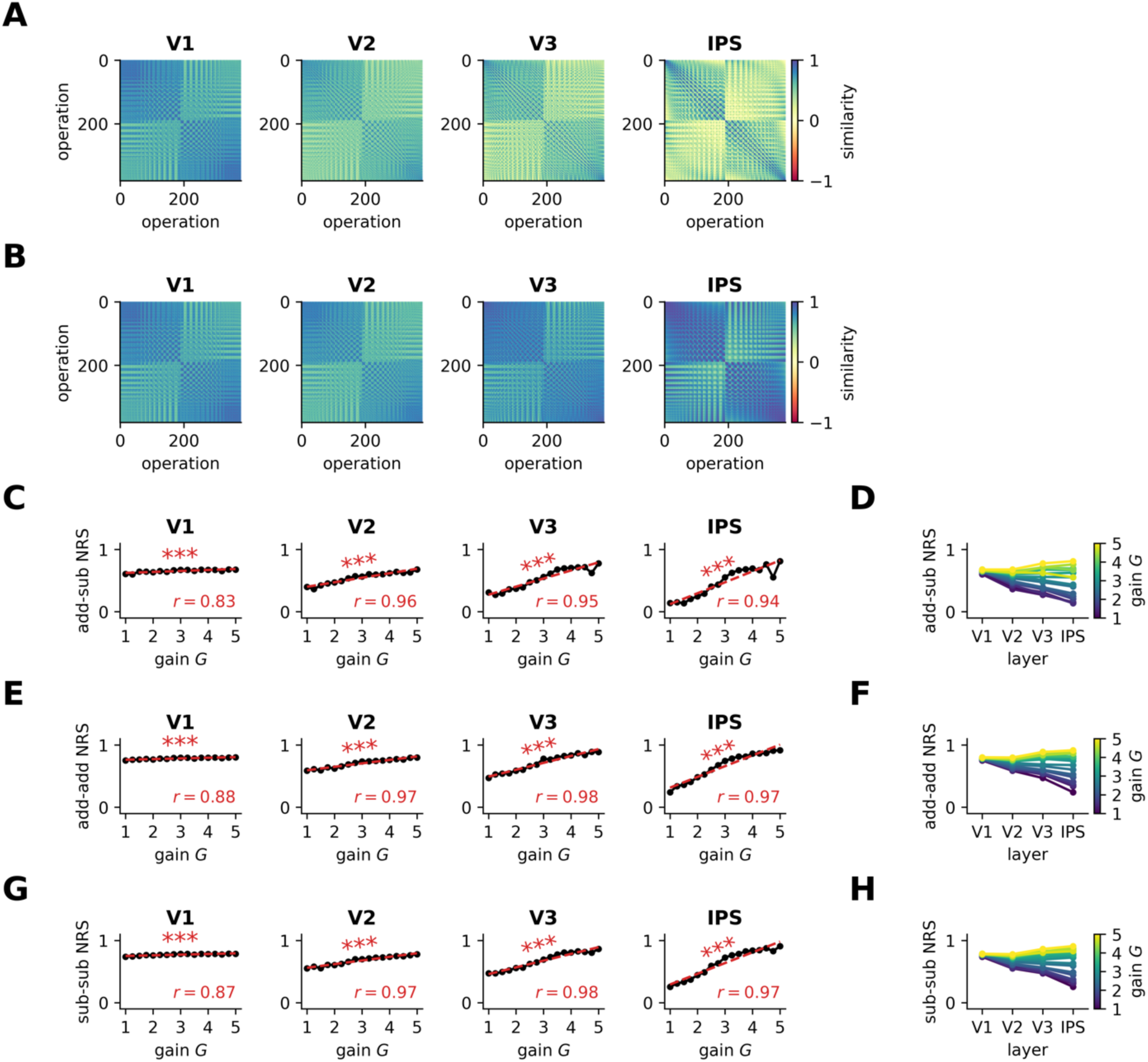
Hyper-excitability diminishes differentiation of neural representations across pDNN layers. **A.** Observed pDNN NRS matrix across model layers (V1→IPS) for a low level of neural excitability (*G* = 2.25). **B.** Observed pDNN NRS matrix across model layers (V1→IPS) for a high level of neural excitability (*G* = 4). **C-D.** Relationship between pDNN NRS and neural gain *G* for NRS between addition and subtraction problems, with linear regression lines (red) showing the strength and direction of the correlation. Summary showing the evolution of NRS measures across model layers (V1→IPS) for different levels of neural gain *G*, depicted as a color gradient from blue (*G* = 1) to yellow (*G* = 5). NRS between addition and subtraction problems (add-sub NRS) increases with neural gain across all layers, indicating reduced differentiation between problem types. **E-F.** NRS between addition problems (add-add NRS) increases with neural gain across all layers, suggesting reduced differentiation within problem types. **G-H.** NRS between subtraction problems (sub-sub NRS) also increases with neural gain across all layers, further confirming reduced differentiation within problem types. *** *p* < 0.001.

To understand how neural excitability influenced NRS, we obtain the average NRS between addition and subtraction problems (*add-sub NRS).* Our findings revealed that after an initial decrease in NRS upon training, the NRS decreased only slightly with further training iterations across layers, even when performance accuracy increased to 95% levels (**Figure SI 1A**). We observed a consistent increase in NRS with neural excitability across all layers (Figure 4C), with a strong correlation between add-sub NRS and neural excitability at each layer: V1 (*r* = 0.83, *p* < 10^−4^), V2 (*r* = 0.96, *p* < 10^−9^), V3 (*r* = 0.95, *p* < 10^−8^) and IPS (*r* = 0.94, *p* < 10^−7^). Figure 4D shows that for low (high) levels of neural excitability, the NRS values decrease (increase) across layers V1 to IPS, and that increasing excitability causes NRS to increase faster in higher layers (V3, IPS) compared to V1 and V2.

Similarly, we examined the NRS between distinct addition problems (*add-add NRS*) and the NRS between distinct subtraction problems (*sub-sub NRS*) and observed a consistent increase in NRS with neural excitability across all layers (Figures 4E-H): *add-add NRS* V1 (*r* = 0.88, *p* < 10^−5^), V2 (*r* = 0.97, *p* < 10^−10^), V3 (*r* = 0.98, *p* < 10^−11^) and IPS (*r* = 0.97, *p* < 10^−10^), and *sub-sub NRS* V1 (*r* = 0.87, *p* < 10^−5^), V2 (*r* = 0.97, *p* < 10^−10^), V3 (*r* = 0.98, *p* < 10^−10^) and IPS (*r* = 0.97, *p* < 10^−9^).

These results demonstrate that increasing neural excitability is sufficient to cause greater neural representational similarity between problem types, that this increased similarity is more pronounced in the higher IPS layer, and that the higher NRS is not completely mitigated with additional training iterations, even when behavioral performance improves to 95% accuracy levels.

### pDNN models of behavioral patterns predict neural task fMRI data reflecting differences between TD and MLD

Extending our analysis to pDNN models representative of children with MLD and their TD peers, we discovered elevated add-sub NRS in MLD-associated pDNN models (Figure 5A) across layers. A statistically significant higher add-sub NRS was noted for MLD pDNNs compared to TD pDNNs at all layers: V1 (*t* = 6.86, *p* < 10^−7^), V2 (*t* = 9.26, *p* < 10^−11^), V3 (*t* = 11.78, *p* < 10^−14^), and IPS (*t* = 12.11, *p* < 10^−14^). Notably, the effect sizes, as measured by Cohen’s d, of the difference in NRS between MLD and TD are large and increase along the network hierarchy: *d* = 2.08 in V1, *d* = 2.82 in V2, *d* = 3.53 in V3, and *d* = 3.66 in IPS.

**Figure 5.**
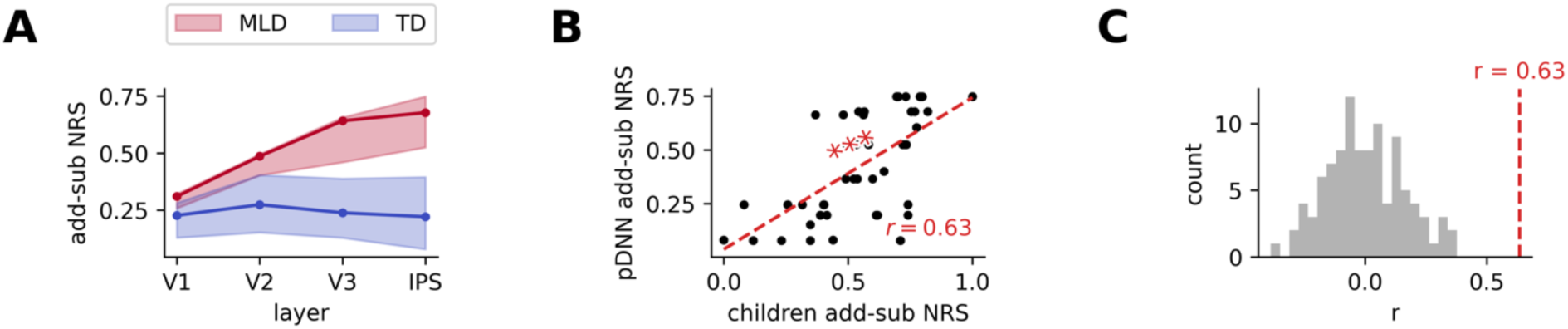
Digital twins predict children’s neural differentiation deficits and validate the excitability-based mechanism underlying math learning disabilities (MLD). **A.** Comparison of NRS between addition and subtraction problems (add-sub NRS) across pDNN layers (V1 to IPS) for models matched to children with MLD (red) and typically developing (TD) children (blue). MLD pDNNs show significantly higher add-sub NRS compared to TD pDNNs across all layers, indicating reduced neural differentiation between problem types. The effect size of the difference in add-sub NRS between MLD and TD pDNNs increases along the network hierarchy, suggesting a more pronounced deficit in higher-order processing regions. **B.** Correlation between children’s empirically observed add-sub NRS in the intraparietal sulcus (IPS), based on brain imaging data, and the predicted add-sub NRS from their corresponding digital twin (behaviorally matched pDNNs). The significant positive correlation (*p < 1e-5*) demonstrates that pDNNs capture the individual variability in neural differentiation deficits observed in children. **C.** Comparison of the correlation between predicted and observed add-sub NRS for behaviorally matched pDNNs (blue) and randomly matched pDNNs (gray). The correlation for behaviorally matched pDNNs is significantly higher than that of randomly matched pDNNs, validating the importance of aligning neural excitability levels to individual behavioral profiles for predicting neural deficits. These results support the neural validity of the pDNN models as digital twins and highlight the critical role of neural excitability in shaping the neural representational deficits observed in children with MLD. *** *p* < 0.001.

Supplementary analysis (**Figures SI 2B-C**) shows that MLD pDNNs also showed higher levels of within operation NRS (both *add-add NRS* and *sub-sub NRS*) compared to TD pDNNs: V1 (*t* > 7.34, *p* < 10^−8^), V2 (*t* > 10.35, *p* < 10^−12^), V3 (*t* > 11.75, *p* < 10^−14^), and IPS (*t* > 10.79, *p* < 10^−13^).

An important aspect of our study was to determine if the optimally tuned pDNNs could reasonably predict NRS deficits in children with MLD, as observed through task fMRI data ^21^. Specifically, we assess whether the pDNNs, calibrated and matched based only on individual task-distal mathematical achievement levels of children, also mirrored the unique NRS patterns evident in the task-related fMRI data from the same children. To achieve this, we calculated the predicted addition-subtraction (add-sub) NRS within the IPS node of the pDNN model that was a best fit to each child’s behavioral data. This prediction was then compared to the actual add-sub NRS derived from previously published empirical fMRI data focusing on the IPS ^21^.

Figure 5B compares the add-sub IPS NRS between the pDNNs and corresponding children. Our analysis revealed a moderately strong positive correlation between the predicted and observed add-sub NRS (*r* = 0.63, *p* < 10^−5^), significantly surpassing the correlation levels that would be expected from random matching (Figure 5C, *t* = 39.70, *p* < 10^−61^).

These results align with our previous empirical findings ^21^ and further demonstrate a clear relationship between increasing neural excitability and heightened representational similarity across numerical operations within the pDNN. This trend is particularly evident in the IPS, suggesting that hyperexcitability in neural networks may underpin the observed phenomenon.

These results underscore an important facet of our pDNN models: their ability to not only align with behavioral patterns but also reflect neural processing differences in the brain. Models that closely mirrored children’s behavioral performance also showed a higher fidelity in approximating neural representation patterns. This alignment provides a compelling validation for the use of pDNNs as a reliable tool in modeling both the behavioral and neurophysiological aspects of mathematical learning and difficulties.

### Neural hyper-excitability impairs both numerical precision and trueness of response by reducing the number of different responses accessible to the network

Behavioral deficits observed in MLD pDNNs may be caused by either a lack of trueness (i.e. average response is far from the true response) or a lack of precision (i.e. response is highly variable around the true value) ^61^. Figure 6A shows the distribution of pDNN responses for each value of the true result for different values of neural excitability at iteration 800. Using these distributions, we compute the numerical systematic error (lack of trueness) as the average distance between the expected response and the average response for that expected response, and we compute the numerical imprecision as the average standard deviation of the response for each expected response. We hypothesized that for pDNNs, both systematic error and imprecision would decrease with training and increase with neural excitability to reflect variations in accuracy.

**Figure 6.**
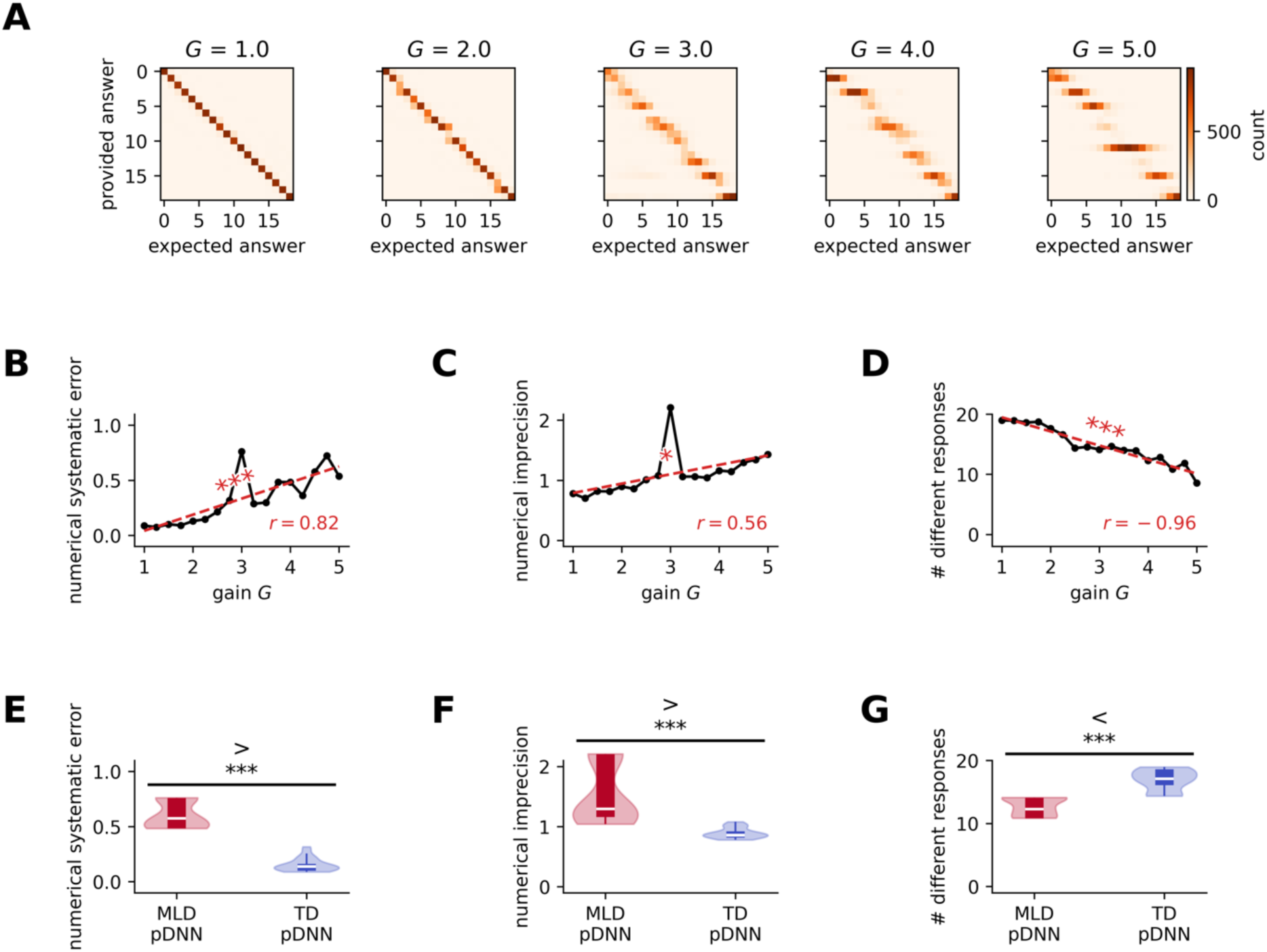
Neural hyper-excitability impairs precision of response and increases systematic error in pDNN models. **A.** Increasing neural gain (excitability) yields more diffuse pDNN response distributions across possible solutions, indicating declining precision. **B.** Numerical systematic error increases with excitability. **C.** Numerical imprecision, measured by standard deviation of responses, increases with excitability. **D.** The effective number of unique responses used by pDNNs, estimated using the entropy of the response distribution, decreases with increasing excitability. This suggests that higher excitability leads to a less diverse set of responses, potentially indicating a less precise internal representation of the numerical solution space. **E.** Comparing the behaviorally matched digital twins for TD and MLD, the numerical systematic error is significantly higher (*p < 1e-14*) for MLD pDNNs compared to TD pDNNs. **F.** Numerical imprecision is significantly higher (*p < 1e-9*) for MLD pDNNs compared to TD pDNNs, aligning with empirical behavioral deficits observed in children with MLD. **G.** The MLD pDNN uses significantly fewer unique responses compared to the TD pDNN (*p < 1e-10*), further supporting the notion that hyper-excitability in MLD is associated with a less precise and less diverse internal representation of numerical quantities. These results highlight the impact of neural excitability on the precision and variability of behavioral responses in pDNNs. * *p* < 0.05, *** *p* < 0.001.

As expected, Figure 6B shows that at iteration 800 the numerical systematic error increases with neural excitability (*r* = 0.82, *p* = 10^−4^), indicating response that are less true. Figure 6C shows that numerical imprecision also increases with neural excitability (*r* = 0.56, *p* = 0.019), indicating that responses are more variable and less precise.

Moreover, as neural excitability increases from 1 to 6, the pDNN responses become both less true and less precise because they are clustered around a fewer number of unique values. First on a qualitative level, we can observe in Figure 6A that for pDNNs, the number of different responses used to provide an answer seems to decrease with neural excitability (fragmentation along the diagonal as gain increases). By estimating the effective number of different responses using the entropy of the response (see details in **Methods**), we note that while pDNN accuracy across training iterations is strongly negatively correlated with both numerical systematic error (*r* = −0.69, *p* < 10^−95^) and numerical imprecision (*r* = −0.73, *p* < 10^−111^), it is even more strongly correlated with the effective number of different responses used (*r* = 0.95, *p* < 10^−323^). Figure 6D shows that the effective number of responses used decrease with neural excitability (*r* = −0.96, *p* < 10^−9^), from using all the 19 unique responses for smaller gains, to only between 8 and 9 responses for higher gains.

This shows how increasing excitability directly translates into behavioral deficits in MLD pDNNs compared to TD pDNNs. Figure 6E shows that the systematic error of MLD pDNNs (*M* = 0.60, *SD* = 0.12) is higher than that of TD pDNNs (*M* = 0.15, *SD* = 0.06) (*t* = 15.40, *p* < 10^−14^). Figure 6F shows that the numerical imprecision of MLD pDNNs (*M* = 1.53, *SD* = 0.49) is higher than that of TD pDNNs (*M* = 0.89, *SD* = 0.09) (*t* = 5.80, *p* < 10^−9^). Figure 6G shows that the effective number of responses used by MLD pDNNs (*M* = 12.71, *SD* = 1.34) is lower than that by TD pDNNs (*M* = 16.89, *SD* = 1.62) (*t* = −9.10, *p* < 10^−10^).

These results suggests that in the pDNNs, the number of results for which there is a precise response increases progressively with training (i.e., learning helps formulate a more precise latent number line) but that increasing neural excitability slows down this learning process, affecting the trueness, precision, and granularity of responses.

### Neural hyper-excitability impairs the structure of representational manifolds

Our next goal was to identify which representational deficits in the pDNNs could cause the behavioral differences observed in MLD vs TD pDNNs. For each layer we studied the geometric properties of the 19-manifold (**Methods**) formed by the neural response to the operations which share the same results (e.g. the response of 2+7 and 5+4 are part of the same manifold as they both result in 9). Specifically we studied their manifold capacities, their dimensionality, and how their manifold centers correlate, as developed in a theory of object maniolds in neural networks ^58,59^. Manifold capacity reflects how easy it is to separate manifold in two random categories, and high (low) manifold capacity indicates that it is easy (hard) to separate the manifolds into two categories. Manifold dimensionality reflects the number of effective dimensions within which the manifold evolves. Correlation between the centers of the manifold reflects the alignement between manifolds, with high correlations indicating that the center of the manifolds are aligned, and low correlations indicating that each center is maximally spread across multiple dimensions. We tested the hypothesis that higher neural excitability should also result in impaired manifold structures. In the context of the current task, lower manifold capacity and higher manifold center correlations reflect impaired manifold structures.

**Figure SI 4A** shows how the manifold capacity evolves during training across layers. After a few iterations we observed, as per our hypothesis, a consistent decrease in manifold capacity with neural excitability across all layers and iterations. At iteration 800 we observed a strong negative correlation between manifold capacity and neural excitability at each layer: V1 (*r* = −0.77, *p* < 10^−3^), V2 (*r* = −0.76, *p* < 10^−3^), V3 (*r* = −0.80, *p* < 10^−3^) and IPS (*r* = −0.76, *p* < 10^−3^, Figure 7A). **Figure SI 5A** shows that at iteration 800, as observed in previous studies ^59^, the manifold capacity increases across layers V1 to IPS, with smaller increases at high levels of excitability.

**Figure 7.**
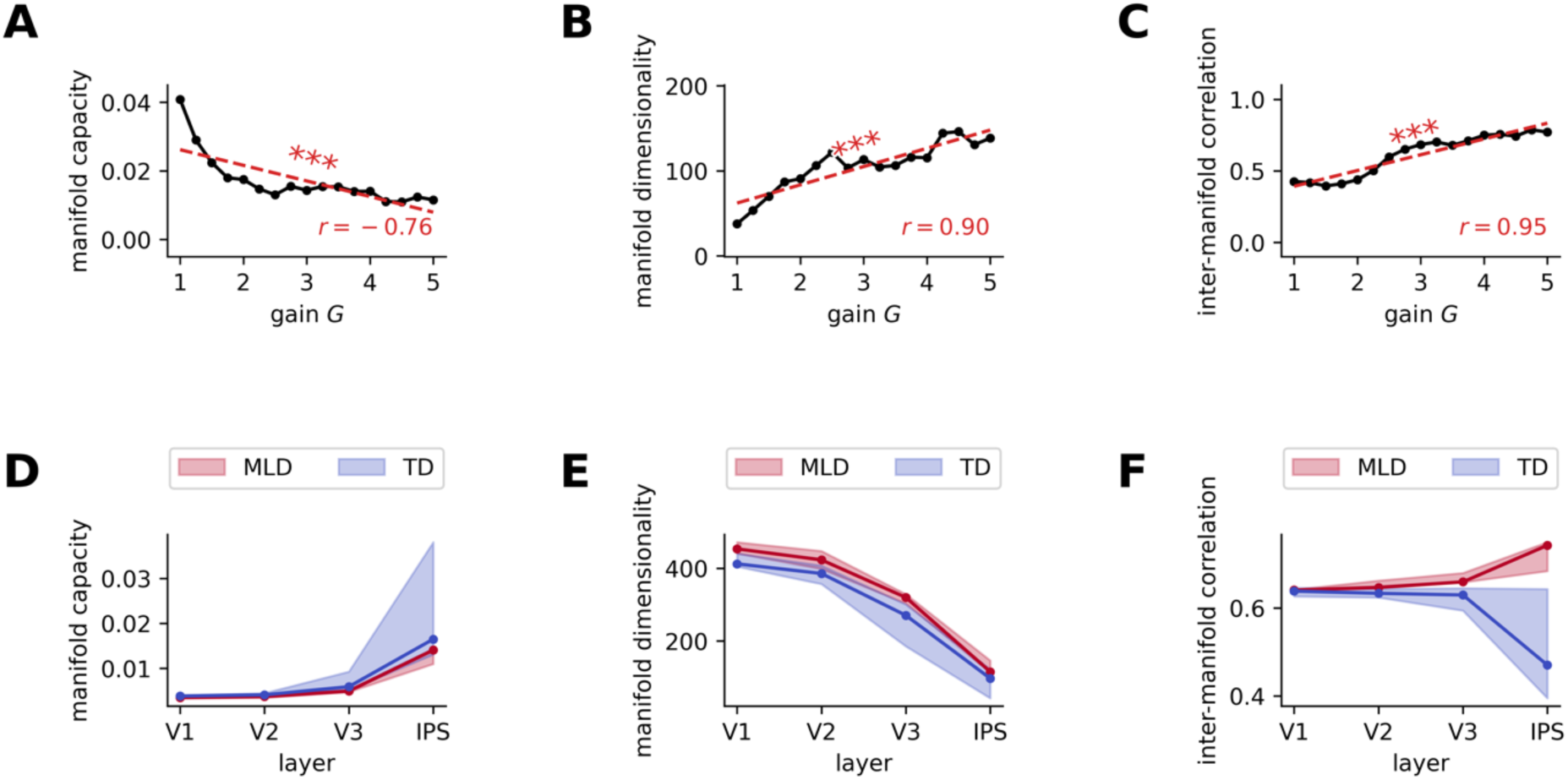
Neural hyper-excitability degrades manifold geometry of latent representations in the pDNNs. **A-C.** Three key manifold properties in the IPS layer of pDNNs change with neural gain levels. **A.** Manifold capacity, reflecting the separability of neural representations, shows a decrease with higher excitation, indicating that hyper-excitability makes it more difficult to distinguish between different numerical manifolds. **B.** Manifold dimensionality, indicating the complexity of the representational space, increases with greater neural gain, suggesting that hyper-excitability leads to more complex and less efficiently organized representations. **C.** Correlations between manifold centers, relating to the alignment of representations, increase with neural gain, implying that hyper-excitability causes the centers of different numerical manifolds to become more aligned, potentially leading to increased interference between representations. **D-F.** At the best-fit iteration, MLD pDNNs (red) exhibit properties consistent with hyper-excitation compared to TD pDNNs (blue). **D.** Manifold capacity is significantly reduced in MLD pDNNs, indicating less separable and more overlapping representations. **E.** Manifold dimensionality is significantly higher in MLD pDNNs, suggesting more complex and less efficient representational spaces. **F.** Correlations between manifold centers are significantly higher in MLD pDNNs, implying suboptimal representations. These results demonstrate that neural hyper-excitability, as observed in MLD, leads to degraded manifold geometry of latent representations in pDNNs. *** *p* < 0.001.

**Figure SI 4B** shows how the manifolds dimensionality evolves during training across layers. After a few iterations we observed a consistent increase in the manifold dimensionality with neural excitability across all layers and iterations. At iteration 800 we observed a strong correlation between manifold dimensionality and neural excitability at each layer: V1 (*r* = 0.93, *p* < 10^−4^), V2 (*r* = 0.80, *p* < 10^−3^), V3 (*r* = 0.87, *p* < 10^−5^) and IPS (*r* = 0.90, *p* < 10^−6^, Figure 7B). **Figure SI 5B** shows that, as observed in previous studies^59^, the manifold dimensionality decreases across layers V1 to IPS.

**Figure SI 4C** shows how the correlation between centers of manifolds evolve during training across layers. We observe a consistent increase across iterations in this correlation in layer V3 and IPS but not in layers V1 and V2. At iteration 800 we observed lower correlations between neural excitability and manifold center correlations in V1 (*r* = 0.005, *p* = 0.99) and V2 (*r* = 0.24, *p* = 0.36), but higher correlations in V3 (*r* = 0.70, *p* < 10^−2^) and IPS (*r* = 0.95, *p* < 10^−8^, Figure 7C). The correlation between manifold centers in the IPS are a strong predictor of behavioral accuracy (*R*^2^ = 0.94). **Figure SI 5C** shows that for low levels of neural excitability, the correlation between centers decreases across layers V1 to IPS, but that for high level of neural excitability, the correlation between centers increases across layers V1 to IPS.

Extending our analysis to pDNN models representative of children with MLD and their TD peers, we discovered reduced manifold capacity in MLD pDNN models (Figure 7D) across all layers. There was a progressive increase in manifold capacity across the pDNN layer hierarchy in both MLD and TD pDNNs. A statistically significant smaller manifold capacity was noted for MLD pDNNs compared to TD pDNNs at all layers: V1 (*t* = −7.74, *p* < 10^−8^), V2 (*t* = −10.37, *p* < 10^−12^), V3 (*t* = −6.07, *p* < 10^−6^), and IPS (*t* = −7.06, *p* < 10^−7^). Notably, the effect sizes, as measured by Cohen’s d, of the difference in manifold capacity between MLD and TD are large across the network hierarchy: *d* = 2.34 in V1, *d* = 1.82 in V2, *d* = 1.54 in V3, and *d* = 1.00 in IPS.

We discovered increased manifold dimensionality in MLD pDNN models (Figure 7E) compared to TD pDNNs, across layers. There was a progressive decrease in manifold dimension across the pDNN layer hierarchy in both MLD and TD pDNNs. A statistically significant higher manifold dimensionality was noted for MLD pDNNs compared to TD pDNNs at all layers: V1 (*t* = 8.89, *p* < 10^−10^), V2 (*t* = 11.37, *p* < 10^−13^), V3 (*t* = 6.82, *p* < 10^−7^), and IPS (*t* = 8.03, *p* < 10^−10^). Notably, the effect sizes, as measured by Cohen’s d, of the difference in manifold dimensionality between MLD and TD are large across the network hierarchy: *d* = −2.65 in V1, *d* = 2.05 in V2, *d* = −2.33 in V3, and *d* = −1.56 in IPS.

We also discovered increased center correlations across manifolds in MLD pDNNs compared to TD pDNNs (Figure 7F) across layers. Interestingly, there was a progressive increase in center correlations across the pDNN layer hierarchy in MLD pDNNs, but a progressive decrease across the layer hierarchy in TD pDNNs. A statistically significant higher correlation between center of manifolds was noted for MLD pDNNs compared to TD pDNNs only in IPS: V1 (*t* = −1.45, *p* = 0.15), V2 (*t* = −0.76, *p* = 0.45), V3 (*t* = 0.43, *p* = 0.67), and IPS (*t* = 12.44, *p* < 10^−15^). Notably, the effect sizes, as measured by Cohen’s d, of the difference in manifold center correlations between MLD and TD are large from V2 to IPS: *d* = −0.44 in V1, *d* = −2.56 in V2, *d* = −2.66 in V3, and *d* = −3.45 in IPS.

Finally, we observed that in the IPS, manifold capacity, manifold dimensionality and inter-manifold correlations between centers were all strongly correlated with accuracy. Surprisingly, inter-manifold correlations between centers showed the strongest correlations with behavioral accuracy (*r* = −0.97, *p* < 10^−323^), followed by manifold dimensionality (*r* = −0.92, *p* < 10^−262^), and manifold capacity (*r* = 0.73, *p* < 10^−109^).

These results show that differences on account of higher neural excitability were explained by decreasing manifold capacity, increasing manifold dimensionality, and increasing center correlations between IPS manifolds.

### Additional pDNN training to overcome behavioral deficits in MLD

Our next goal was to use pDNNs to determine how additional training could help MLD pDNNs reach the level of behavioral performance seen in TD pDNNs. We hypothesized that both behavioral and neural representational deficits would reduce with additional cognitive training, and that pDNNs corresponding to MLD children would converge towards behavioral and neural representational patterns observed in TD children.

Figure 8A shows how many additional iterations were required for any individual pDNN to reach an accuracy level closest to the median accuracy of best matching TD pDNNs (i.e., at iteration 800). The number of iterations required to match TD pDNNs increased consistently with neural excitability (*r* = 0.93, *p* < 10^−7^). Figure 8B shows the distribution of percentage of initial training that is additionally required (i.e., the number of additional training iterations required beyond the initial 800, divided by 800) to reach median level accuracy of TD pDNN, separately for MLD pDNNs and TD pDNNs. Expectedly, we observe that MLD pDNNs require a higher level of additional training (*M* = 75%, *SD* = 42%) than TD pDNNs (*M* = 5%, *SD* = 11%). MLD pDNNs required an additional 900 iterations beyond the initial 800, or in other words, MLD pDNNs required approximately 2 times the training given to TD pDNNs to achieve similar levels of accuracy.

**Figure 8.**
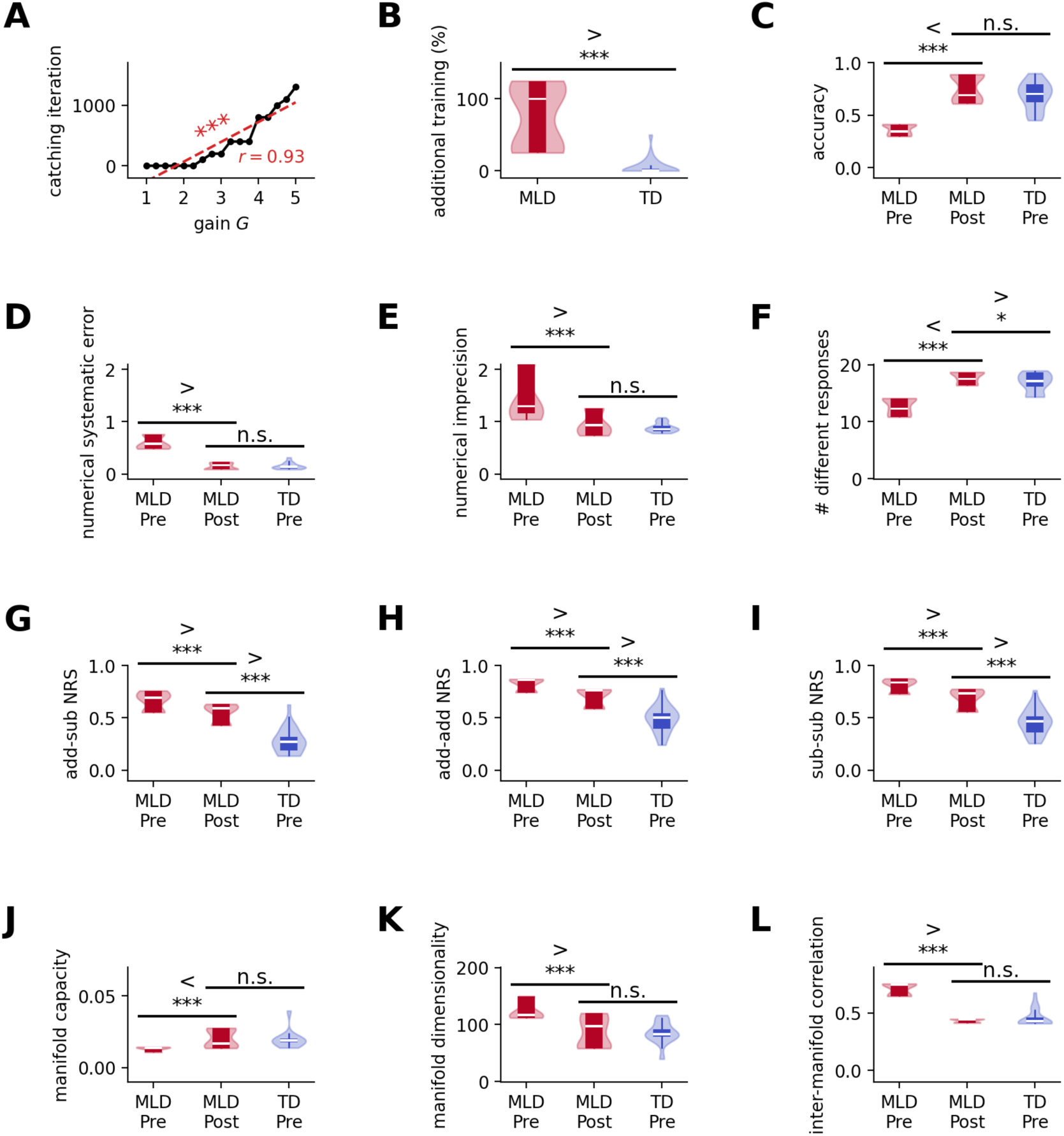
Additional training enables MLD digital twins to remediate behavioral but not all neural representational deficits. **A.** The number of additional training iterations required for MLD pDNNs to catch up to the accuracy level that TD pDNNs demonstrate at iteration 600 increases consistently with higher neural gain. **B.** MLD pDNNs require on average 2.7x the training to reach the same performance level as TD pDNNs. **C-L.** Comparison of MLD pDNN properties pre-training (iteration 800, best behavioral fit) versus post-additional training (iteration 1700, performance caught up to TD levels). **C-F.** Properties of behavioral responses. **G-I**. Similarity between neural representation (NRS). **J-L** Manifolds geometrical properties. **C.** Deficits in accuracy, **D.** numerical systematic error, **E.** numerical imprecision, and **F.** effective number of unique responses are remediated by additional training. However, deficits in NRS persist **G.** across operations (add-sub NRS), **H.** across addition trials (add-add NRS) and **I.** across subtraction trials (sub-sub NRS). Deficits in **J.** manifold capacity, **K**. manifold dimensionality, and **L.** manifold center correlations normalize with additional training. These results suggest that while behavioral deficits can be remediated through additional training, neural representational deficits persists in MLD pDNNs, but aberrant geometry of the underlying manifolds normalize. * *p* < 0.05. *** *p* < 0.001.

We then analyzed how behavior and representations evolved with this behavioral catch-up for MLD pDNNs at iteration 1700, and how they compared to the behavior and representations of TD pDNNs at iteration 800. In Figure 8C-L, we refer to MLD/TD pDNN at iteration 800 as MLD/TD Pre, and MLD pDNN at iteration 1700 as MLD Post.

Figure 8C shows that the accuracies of MLD pDNNs at iteration 1700 improve (*t* = 13.50, *p* < 10^−12^) and are not significantly different from TD pDNNs at iteration 800 (*t* = 1.90, *p* = 0.065). Similar patterns are seen in the numerical systematic error (Figure 8D; significant improvement in MLD pDNNs at iteration 1700, *t* = −15.35, *p* < 10^−14^; not significantly different from TD pDNNs at iteration 800, *t* = −0.17, *p* = 0.86), numerical imprecision (Figure 8E; significant improvement in MLD pDNNs at iteration 1700, *t* = 4.89, *p* < 10^−4^; not significantly different from TD pDNNs at iteration 800, *t* = 1.21, *p* = 0.24), manifold capacity in the IPS (Figure 8J; significant improvement in MLD pDNNs at iteration 1700, t = 5.42, p < 10^−6^; not significantly different from TD pDNNs at iteration 800, t = 0.26, p = 0.79), manifold dimensionality in the IPS (Figure 8K; significant improvement in MLD pDNNs at iteration 1700, t = −5.83, p < 10^−5^; not significantly different from TD pDNNs at iteration 800, t = 0.21, p = 0.84), and inter-manifold center correlations in the IPS (Figure 8L; significant improvement in MLD pDNNs at iteration 1700, t = −24.53, p < 10^−17^; not significantly different from TD pDNNs at iteration 800, t = −1.66, p = 0.11).

Number of different responses (Figure 8E) in MLD pDNNs shows significant improvement with additional training (*t* = 13.26, *p* < 10^−14^) and improves beyond the levels observed in TD pDNNs at iteration 800 (*t* = 2.15, *p* = 0.038).

The remaining measures for MLD pDNNs show an improvement with the additional training to 1700 iterations, but do not statistically reach the levels demonstrated by TD pDNNs at iteration 800. This includes between operation add-sub NRS (Figure 8G; significant improvement at iteration 1700, t = −4.39, p < 10^−4^; still significantly different from TD pDNNs at iteration 800, t = 8.54, p < 10^−9^), within addition NRS (Figure 8H; significant improvement at iteration 1700, t = −5.39, p < 10^−5^; still significantly different from TD pDNNs at iteration 800, t = t = 6.48, p < 10^−7^), and within subtraction NRS (Figure 8I; significant improvement at iteration 1700, t = −5.19, p < 10^−5^; still significantly different from TD pDNNs at iteration 800, t = 6.65, p < 10^−7^).

These results suggests that when MLD pDNNs are provided additional training they can catch up behaviorally to performance levels characterizing TD pDNNs. This behavioral catch-up is accompanied by improvements in all measures of neural representations examined here, but only some of the underlying neural representational deficits are completely overcome, while others improve but still do not catch up to TD pDNN levels.

## Discussion

Our study integrates distinct lines of research in human cognitive neuroscience and artificial intelligence and provides new insights into latent brain mechanisms that contribute to the diversity of children’s cognitive abilities. We developed cutting-edge personalized deep neural networks (pDNNs), which are tailored to probe the neural gains, learning dynamics, and neurophysiological patterns unique to each child (Figure 1), thus providing *in silico* digital twins. These digital twins capture the behavioral, learning, and neural variability demonstrated by both TD children, and children with MLD. We demonstrate that neural network models when combined with individual human behavioral and brain imaging measures offer powerful tools for advancing our knowledge of the neurobiology of cognitive strengths and challenges. Our findings highlight the transformative capacity of pDNNs as a tool in the assessment and exploration of cognitive disabilities, paving the way for more personalized and effective educational strategies.

### Engineering pDNNs as digital twins to probe individual learning profiles

We developed cognitive neuroscience-informed and biologically plausible pDNNs ^14^ to capture individual differences in arithmetic task performance among children with and without MLD. This model was designed to investigate the complex interplay between behavior, neural representations, and neurophysiology in children with learning disabilities. We engineered pDNN models with the unique ability to map diverse arithmetic problems to corresponding solutions, effectively tuning these models to align with various performance levels. This allowed us to investigate the latent neurobiological mechanisms that might contribute to the observed individual differences and weaknesses in neural representations during problem solving.

We demonstrate that these pDNNs can be tuned to match individual levels of mathematical achievement across varying levels of performance abilities, and that such a model can be used to uncover latent neurobiological mechanisms underlying weak neural representations of distinct numerical problems. A key theoretical aspect of our work is analysis of the role of excitatory-inhibitory (E/I) imbalances^37,39,43,55,56^, manifested as neural hyper-excitability, in capturing the variability in behavioral, neural, and representational dysfunctions observed in children with MLD.

Our methodology encompassed six key steps: First, we employed a biologically inspired hierarchical pDNN model, mimicking the dorsal visual pathway from V1 to IPS, to encode addition and subtraction problems. This model’s design was based on the CORNET-S architecture, similar to one used to model the emergence of number sense in children ^14^. Second, we adjusted the neuronal excitability parameters within these models to induce individual differences in task performance. Third, data from children participating in the empirical study were matched with a corresponding pDNN configuration that best represented their behavioral performance, to obtain *in silico* digital twins corresponding to each child. Fourth, we then explored the congruence in behavioral performance and in neural representations between the pDNNs and empirical task-based fMRI studies. Fifth, we investigated how variations in neural hyper-excitability impacted the latent neural representations within these networks, aiming to shed light on potential neurophysiological underpinnings of MLD. Finally, we probed the influence of additional training on pDNN networks in MLD, and determined whether latent neural representations were normalized by additional cognitive training. The resulting engineered ‘digital twin’ models provide a novel approach to close the loop between AI-based neural networks, behavior, neural representations, and neurophysiology.

### Neural excitability impairs learning

Our study brings to light the significant impact of neural excitability and hyper-excitability on learning processes, which we evaluated here in the context of MLD. Theoretical models propose that E/I imbalances could disrupt functional neural circuitry, potentially leading to a range of neurodevelopmental disorders, including MLD ^41,43,56,57^. By focusing on E/I imbalance, our pDNN model provides a more precise understanding of the neurobiological underpinnings of learning disabilities. This approach also offers a new perspective for investigating the complex interplay between neural circuitry and learning processes ^41,43^.

A central finding is how neural hyper-excitability influences the capacity to learn associations between mental arithmetic problems and their solutions. We observed that elevated neural excitability levels in the pDNNs were correlated with slower learning rates and less effective learning outcomes (Figure 2). The results further revealed that as neural excitability increased, the learning rate of the pDNNs decreased in a monotonic fashion. This trend was not just a delay in the learning curve but also a qualitative alteration in the learning dynamics of the pDNNs associated with poor performance.

Next, we fine-tuned pDNNs to match the behavioral performance of individual children by adjusting neural excitability values. The behavioral scores of matched pDNNs closely mirrored the empirical behavioral patterns observed in TD and MLD children, validating the utility of pDNNs in modeling the behavioral nuances of children with MLD and TD in mental arithmetic tasks. The neural excitability (gain value *G*) in MLD pDNNs was found to be significantly higher than in TD pDNNs. This indicates that neural excitability plays a crucial role in personalizing pDNNs to represent varying levels of task performance, with MLD pDNNs associated with higher neural excitability (Figure 3). Thus, neural excitability maybe a significant contributor to suboptimal performance levels observed in children with MLD. These results resonate with empirical research on Glutamate/GABA levels indicating excitatory-inhibitory imbalances ^41,43,56,57^, and intrinsic fMRI hyperactivity and hyperconnectivity ^34^, as well as task-related hyperactivity and hyperconnectivity in MLD ^18^.

Together, our computational modeling of pDNNs and empirical findings reinforce the notion that neural excitability plays a crucial role in learning and may be a key neurophysiological factor underlying MLD. By highlighting the role of neural hyper-excitability, our research advances understanding of the etiology and manifestation of MLD. Furthermore, it points to neural excitability as a promising biomarker for identifying learning challenges, guiding interventions, and monitoring their effectiveness.

### Neural hyper-excitability leads to less differentiated neural representations in pDNNs, and predicts empirical fMRI findings

Next, we examined the impact of neuronal hyper-excitability on distributed neural representations in each layer of the pDNN model. Neural representational similarity (NRS) was used to quantify the degree of overlap or distinctiveness between neural representations of addition and subtraction problems^62^. NRS measures the correlation between patterns of neural activity evoked by different stimuli or task conditions. High NRS indicates that the neural representations are similar or overlapping, while low NRS suggests more distinct or differentiated representations.

We found that increased neuronal excitability led to less differentiated neural representations in processing numerical operations (Figure 4). Specifically, NRS between addition and subtraction problems (add-sub NRS) was consistently higher across all layers of the pDNN as neural excitability increased. This effect was most pronounced in the IPS layer of the model, a key brain region for numerical cognition. Such diminished differentiation in neural representations in the IPS is significant, as it aligns with empirical findings in children with MLD, who also exhibit less distinct neural representations across numerical operations ^21^. It is also notable that the differences in NRS between TD and MLD pDNNs were minimal in the perceptual V1 layer, but were significantly amplified at subsequent cognitive stages, with maximum differences in the IPS.

Furthermore, we examined within-operation NRS across addition problems (add-add NRS) and subtraction problems (sub-sub) and found that it also increased with higher neural excitability across all pDNN layers (Figure 4). This suggests that hyper-excitability not only leads to greater overlap between distinct numerical operations but also blurs the boundaries between representations of problems within the same operation type.

Importantly, by tuning the neural excitability of the pDNN to align with individual behavioral variations in children, we could accurately map variations in neural representations to empirical fMRI data. Specifically, we determined whether neural representations in pDNNs could predict empirical values of neural representations obtained in brain imaging data obtained from the same group of children. Even though the models were fit only on task-distal behavioral achievement levels, we found that the best-fitting pDNN models showed a reasonably strong level of prediction for the severity of aberrancies in neural representations deficits as observed in task-based fMRI data from children with MLD (Figure 5). This alignment between predicted and observed neural representations underscores the relationship between neural excitability and representational similarity across numerical operations within the pDNN.

Together, our findings demonstrate how tuning neural excitability in pDNNs can effectively model individual differences in math abilities, highlighting the role of neural hyper-excitability in cognitive performance deficits among children with MLD. Additionally, the findings emphasize the potential of pDNNs to predict high-level neural representations, offering insights into neurobiological dysfunction associated with learning disabilities.

### Neural excitability reduces the trueness, precision, and granularity of behavioral responses

Next, we examined how neural excitability influences the trueness, precision, and granularity of behavioral responses in pDNNs that simulate the cognitive performance of children with MLD and TD children. Trueness refers to how close, on average, the responses are to the correct answers, and is measured by systematic errors from the true answer. Precision on the other hand refers to how consistent or variable the responses are for a given answer. The granularity is measured in terms of number of different responses that the pDNN is able to provide across the set of problems. We found that higher neural excitability led to higher systematic errors, lower precision (higher variability around the true answer), and lower granularity of responses. (Figure 6B-D). This suggests that neural excitability affects both the accuracy and the consistency of the pDNNs’ performance on the mathematical problem-solving tasks, along with the use of a smaller subset of possible responses. The use of a smaller set of unique responses suggests a less developed internal representation of numerical magnitudes. This further supports the idea that higher neural excitability constrains the pDNNs’ ability to develop a rich and precise representation of the numerical space. These results were mirrored in our comparison of MLD and TD model pDNNs, with higher errors, lower precision, and a smaller number of unique responses in the MLD group compared to the TD group (Figure 6 E-G).

This highlights a plausible neurobiological mechanism for the behavioral deficits observed in children with MLD. The pDNN model provides a framework for understanding how abnormalities in neural excitability can give rise to the cognitive and behavioral impairments associated with MLD. This insight can guide future research on potential interventions that target specific neural mechanisms underlying MLD, such as training paradigms that aim to improve systematic errors, imprecision, and granularity of responses.

### Neural excitability influences manifold structure

Our next objective was to uncover how neural excitability impacts the separability and geometry of object manifolds in pDNNs ^58,59^. Stimuli are represented in the brain by the collective population responses of neurons, and an object presented under varying conditions gives rise to a collection of neural population responses called an ‘object manifold’ ^59,63^. Manifolds in this context represent low-dimensional subspaces formed by neural responses to stimuli that share common features. To explore the geometric properties of neural responses we leveraged recent advances in manifold analysis, focusing on three key measures: manifold capacity, manifold dimensionality, and inter-manifold center correlation. Recent theoretical progress has connected these measures with classification capacity ^58,59^. By examining alterations in the manifold structure of latent representations across various layers of the pDNN, we aimed to quantify how neural network properties are altered by neural gain and elucidate the impact on pDNN models associated with MLD.

Manifold capacity reflects the ease of separating a collection of manifolds into two categories. A high manifold capacity signifies greater ease in segregating the manifold into two discernible categories. In the present study, categories were represented by the 19 possible solutions ranging from 0 to 18 as described above. As expected, we found that manifold capacity increased with training (**Figure SI 4**). Crucially, we found consistent decrease in manifold capacity with increased neural excitability, demonstrating that higher neural excitability makes it harder to distinguish between different numerical representations (Figure 7A). Analysis comparing pDNN models in MLD and TD revealed that manifold capacity increased progressively from layers from V1 to IPS, with higher capacity in TD compared to MLD pDNN models (Figure 7D), and lower amplification of the manifold capacity as we move across the stages in the cognitive hierarchy from V1 to IPS in MLD pDNNs. This differentiation underscores the impact of neural excitability on numerical cognition, highlighting a critical barrier in MLD that hampers the separation of numerical representations into clear, distinguishable categories.

Manifold dimensionality denotes the requisite number of effective dimensions to encapsulate the geometric characteristics inherent to the dataset. We observed a significant increase in manifold dimensionality with neural excitability (Figure 7B). This suggests that elevated neural excitability creates a more complex and less easily partitioned representational space, complicating the task of distinguishing between distinct problem solutions. Analysis comparing pDNN models in MLD and TD revealed a progressive decrease in manifold dimensionality from the V1 to IPS layers, with lower dimensionality in TD compared to MLD pDNN models (Figure 7E).

Finally, we examine inter-manifold correlation which quantifies the relation between centers of manifolds across the 19 possible solutions. High correlations would indicate that the centers of the manifolds are aligned, while low correlations indicate that each center is maximally spread across multiple dimensions. Inter-manifold correlations in the IPS, but not earlier layers, increased with neural excitability (Figure 7C), suggesting that higher excitability leads to impairments in how the manifolds are organized in the neural space at a higher cognitive (IPS) but not lower perceptual (Figure 7F) level. Interestingly, inter-manifold correlation, rather than manifold capacity or dimensionality, was the most predictive of pDNN performance on both addition and subtraction problems. Analysis comparing pDNN models in MLD and TD revealed a striking difference in the pattern of inter-manifold correlation across layers. In TD pDNNs, inter-manifold correlation decreased progressively from lower (V1) to higher (IPS) layers, suggesting a gradual decorrelation of problem set representations along the processing hierarchy (Figure 7F). This aligns with previous reports of reduced correlation between neural representations in higher processing stages of deep neural networks, which is thought to support more efficient and robust information processing ^59^. In contrast, MLD pDNNs showed a progressive increase in inter-manifold correlation from lower to higher layers, with a higher correlation in the IPS layer compared to TD pDNNs (Figure 7F). This suggests that the neural representations of different problem sets are sub-optimally separated in the higher processing stages of MLD pDNNs, which may contribute to the deficits in numerical problem-solving observed in this group.

These results demonstrate that the manifold structure of latent representations changes across various layers of the pDNN, with the IPS showing highest manifold capacity, and lowest manifold dimension and inter-manifold correlation. Each of these geometric properties was distorted in pDNN models associated with MLD. The findings provide a more comprehensive model of how neural excitability influences not just overall learning performance, but also the internal geometric structure of neural representations. These impediments highlight manifold structure as a potential neural marker for distinguishing between typical and atypical numerical processing pathways and provide clues towards the design of training and interventions that could target specific aspects of neural representations, such as inter-manifold center correlations. Digital twin platforms could provide an experimental setup to test how different training paradigms affect different aspects of manifold geometry.

### Mitigating behavioral deficits in MLD through extended learning

Next, we examined whether extended learning could mitigate behavioral deficits in MLD. Remarkably, we found that with sufficient training MLD pDNNs can reach the same proficiency levels in mathematical tasks as pDNNs tuned to performance levels tuned to TD controls (Figure 8C-E). This implies that children with MLD may require more time and training to achieve the same level of proficiency as TD children. The amount of additional training needed was directly proportional to individual levels of hyper-excitability (Figure 8A). We observed that on an average, MLD pDNNs required about 2 times the training required by TD pDNNs to reach the same average levels of behavioral performance (Figure 8B). While this is a significantly higher level of time and effort, the positive finding is that while neural hyper-excitability significantly slows down learning, it may not be an unsurmountable impediment to learning. Such a delay in learning progression, characterized by a decrease in the learning rate in proportion to increased neural excitability, aligns with empirical evidence suggesting that with targeted and sustained training, children with MLD can progressively improve arithmetic task performance ^64^. This finding suggests potential pathways for intervention that could help children with MLD achieve their full learning potential. Future studies employing rigorous cognitive training methodologies are needed to validate the potential of such interventions. These studies should aim to explore the optimal intensity, duration, and type of cognitive training that would be most beneficial for children with MLD.

### Persistent MLD deficits in latent neural representations and manifold structure

Our investigation next focused on whether additional training, which normalized behavioral performance in MLD pDNNs to match that of TD pDNNs, also normalized latent neural representations as assessed by neural representational similarity and object manifold properties. Despite a 2-fold increase in training for MLD pDNNs, we observed that improvements in behavioral performance did not correspond to equivalent changes in all aspects of neural representations and manifold structures.

While manifold geometrical properties, such as manifold capacity, manifold dimensionality and center correlations of the manifold structures significantly, changed for the MLD pDNNs with additional training (Figure 8L), aligning with the levels shown by the best-matched TD pDNNs, other deficits persisted. Specifically, although neural representational similarity decreased with additional training, MLD pDNNs continued to exhibit higher neural representational similarity compared to TD pDNNs (Figure 8G-I). This suggests that certain latent neural representations remain resistant to change even after extensive training. This selective influence on latent neural representations highlights specific mechanisms that could be targeted for remedial cognitive interventions, although the precise approaches require further investigation.

### Implications for educational neuroscience

Our findings have implications for educational neuroscience. The ability to create digital twins that model individual learning processes and neural patterns opens new avenues for personalized education strategies. These strategies could be specifically tailored to address the unique cognitive needs and challenges of each child, particularly those with MLD. The insights gained from our pDNN models suggest the potential for developing more effective intervention strategies for children with MLD. Specific AI based strategies can be used to discover the most effective training paradigms – for instance, evaluating training paradigms that are the most effective in reducing aberrant neural representations and manifold structure could lead to identifying the most effective training mechanisms for addressing learning disabilities.

The fact that neural representations and object manifold structure between problem types were not fully remediated for MLD pDNNs, despite behavioral accuracy normalization, suggests that high neural excitability may present persistent latent neural representations. These representations could impose learning constraints on more complex problem sets or necessitate a significantly higher level of training than explored in this study for mitigation.

Our findings also highlight the persistence of certain neural representational deficits even after behavioral performance has been normalized through additional training. This suggests that while we can improve behavioral outcomes, underlying neural representations may require more targeted and possibly intensive interventions. The selective influence of additional training on latent neural representations, such as the decorrelation of problem set representations, reveals specific neurobiological mechanisms that could form potential targets for cognitive interventions. However, the exact nature and implementation of these interventions remain to be explored in future research.

### Limitations and future directions

While our pDNN approach has provided a novel perspective for investigating the neurobiological underpinnings of mathematical difficulties, it focuses primarily on E/I imbalance as a theoretical mechanism, and proves its sufficiency, but not necessity, for characterizing individual differences in learning profiles. Future research should consider other plausible mechanisms that may contribute to learning disabilities. Our approach demonstrates how alternate hypotheses can be systematically evaluated against empirical data using DNN models. While our study presents a significant step forward, it is important to acknowledge the limitations of modeling complex human cognitive processes such as mental arithmetic using DNNs. The scope of our study was confined to an area of mathematical learning, and the applicability of our findings to other cognitive domains remains to be explored. Future research should explore the applicability of pDNNs to other cognitive tasks and learning disabilities.

Given the significant role of neural excitability and hyper-excitability in learning processes, particularly in the context of MLD, an important implication of our research is the potential for brain stimulation techniques designed to suppress neural hyper-excitability as a therapeutic intervention for MLD. Such interventions could target the excitatory-inhibitory balance in key brain hubs, aiming to normalize neural excitability levels and thereby improve learning outcomes for children with MLD. This approach aligns with the broader goal of developing personalized education and intervention strategies based on individual neurobiological profiles.

### Conclusion

Our study represents a significant advance in the integration of cognitive neuroscience and artificial intelligence to unravel the complex neurobiological mechanisms underlying MLD in children. By developing and employing pDNNs as digital twins, we have elucidated the intricate interplay between neural excitability, learning dynamics, and individual neurophysiological patterns that contribute to the diverse cognitive abilities observed in children. Our pDNN models, informed by cognitive neuroscience and tailored to individual learning profiles, also mirror the learning patterns and neural activity observed in children, thereby validating their utility in cognitive neuroscience. The application of pDNNs to model individual learning processes and neural patterns in children with MLD demonstrates the potential of these models in cognitive neuroscience and opens new avenues for the development of targeted educational interventions.

## Methods

### A. Human study protocol and Design

We developed a pDNN to model arithmetic problem-solving tasks performed by children with mathematical learning disabilities (MLD) and typically developing (TD) children during fMRI scanning ^18,21,34^.

#### Participants

Behavioral and neuroimaging data were acquired from 45 children in their 2nd or 3rd grade of schooling (ages 7 to 9). Numerical problem-solving skills of children were assessed using the Numerical Operations (*NumOps*) subtest of the Wechsler Individual Achievement Test 2^nd^ Edition ^60^. 21 children scoring below 90 (i.e. the 25th percentile) on the *NumOps* were classified in the MLD group, while the remaining 24 children formed the TD group. The two groups did not differ on age, full-scale IQ, and reading abilities. All participants had Full-scale IQ scores > 80 (range: 84-128), as assessed by the Wechsler Abbreviated Scale of Intelligence (WASI).

#### Behavioral task

Children were shown equations involving additional or subtraction operations that were a sum or a difference of two small numbers, e.g. “10 + 2 = 13” or “10 − 2 = 8. They were asked to press one of two buttons, the first identifying the equation as correct, e.g. “10 − 2 = 8”, and the second identifying the equation as wrong, e.g. “10 + 2 = 13”. Additional details on the behavioral task conditions are previously published ^18,21,34^.

#### Neural recordings

Each child performed the task during fMRI scanning. Additional details on the fMRI data acquisition, preprocessing, and analysis procedures are previously described ^18,21,34^.

### B. pDNN study protocol and Design

To probe the impact of neural hyperexcitability on mathematical learning and on representations we adapted a math task that has been studied in our lab in children^18,21,34^ to a task that can be solved by an artificial neural network model. We then adapted a biologically inspired artificial neural network model of the visual cortex to solve that task and personalized these biologically inspired networks to match individual differences in the performance of children by varying the neural excitability of the networks. We observed the impact of varying neural excitability on both the behavioral performance and neural representation used to solve that task. Finally, we compared the observed effects of increased neural excitability to the effects observed in children with MLD.

#### Step 1: Adaptation of behavioral addition and subtraction tasks and stimuli

In the original study, asking for the validity of an equation instead of directly asking for the result of sums or differences was a way to simplify the apparatus used to perform the task while the fMRI signal was being recorded. The pDNN task uses similar addition and subtraction problems, where the model has to produce the right answer. We used the MNIST dataset of handwritten digits ^65^ to generate images of human-readable sums and differences of positive integers. We considered only sums and difference which resulted in a value bounded between 0 and 18, i.e. 380 unique problems (190 unique addition and 190 unique subtraction problems). For each problem we generated 100 variants, half were used for training and the other half for testing. In total we used 19000 problems for training and 19000 problems for testing. Since operation symbols are not present in MNIST, we generated synthetic operation symbols “+” and “–” by using the character "1" as a vertical stroke and rotated this character to obtain a horizontal stroke. We represented each problem as *T*_1_*U*_1_*ST*_2_*U*_2_, where *T*_1_ and *U*_1_ (resp. *T*_2_ and *U*_2_) represent respectively the tenth and unit digit of the first (resp. second) operand, and *S* represents the symbol of the operation (+ or −). For single digit operands we consider the tenth digit to be an empty space (an image filled with black). To generate each variant, we randomly selected a visual representation for each character (i.e. a 3x28x28 tensor filled with 0 for an empty space) and concatenated them into a 3x28x140 tensor (See **Figure SI 6**).

#### Step 2: Using pDNNs to solve addition and subtraction problems

The pDNNs model the dorsal visual pathway involved in numerical cognition. The architecture of pDNN is adapted from CORnet-S, a model of the visual pathway. Our adapted pDNN is composed of four layers V1, V2, V3 and IPS, corresponding to key brain regions forming the dorsal visual processing stream, and participating in the processing of numerical information. All representational analysis of pDNN were focused on the last time step of each layer (see **SI** for details of the architecture). For the sake of simplicity and in order to reduce any initial bias, the pDNN was not pretrained on ImageNet or any other stimuli set, that is, only the architecture from CORnet-S was used, and not its weights after training on ImageNet. Moreover, three additional structural modifications were introduced to the network adapted from CORnet-S. First, the output layer was made 19-dimensional, corresponding to the 19 possible answers between 0 and 18. Second, in order to control the excitability of neurons, the parameters governing batch normalization were fixed across training iterations. In pDNN, the batch normalizations enforce the input to the non-linearity, (i.e. the input to the neurons) to have a mean of 0 and a variance of 1 across training iterations. Third, we modified the non-linearity to account for differences in neural excitability of neurons (see Step 3 for details).

We trained pDNNs to solve visually presented addition and subtraction (see Figure 1A) for different levels of excitability using cross entropy loss as the error function, and the Adam optimizer ^66^ with a learning rate of *η* = 0.001. We tested the pDNNs after every 100 batches of 100 problems, i.e. after learning from 10000 examples of addition or subtraction.

#### Step 3: Varying neural excitability in pDNN

In pDNN, the response of a neuron is simplified as follows:

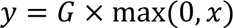

where *y* represents the output firing rate of a neuron, *x* represents the overall input that the neuron receives from other neurons (enforced to be of mean 0 and standard deviation 1 by batch normalization), *G* represents the gain of the neuron that scales the intensity of the response (see Figure 1C). We consider the gains *G* varying from 1 to 5 in steps of 0.25 (Figure 1B), i.e., *G* = 1 + 0.25 × *k* for *k* ∈ ⟦0,16⟧. In the paper, excitability level refers to the gain *G*.

#### Step 4: Behavioral matching

To compare any single pDNN and child at the behavioral level, we calculate a behavioral distance between the normalized accuracy of the pDNN model and the normalized *NumOps* score for the child. These measures are normalized by setting the maximum and minimum accuracy measured over all gains and iterations to 1 and 0 respectively for the pDNNs, and by setting the maximum and minimum *NumOps* scores to 1 and 0 respectively for the children. We then use a Manhattan distance (L1 norm) to define the distance between the normalized *NumOps* score and the normalized pDNNs scores. This is used to find the best matching excitability level of pDNN for each child at each training iteration.

#### Step 5: Identifying the best matching training iteration

We select the best matching iteration as the iteration where the average behavioral distance scores across children and their best-fit models is the smallest. This distance was minimal for iteration 800.

#### Step 6: Behavioral and neural analysis of pDNN

We focused pDNN behavioral analysis on (1) its accuracy on the addition and subtraction problems, and on (2) the trueness, the precision, and the entropy of its response to the problems. Inspired by previous studies (e.g. investigating the differences between TD and MLD children) we examined how neural gain *G* was affecting (1) the neural representation similarity (*NRS*) between addition and subtraction problems within each region of pDNN, and (2) the geometric properties of the 19 manifolds specific to each pDNN response to the problems.

### Numerical systematic error and imprecision

As standardized in ^61^, we defined numerical systematic error and numerical imprecision (represented in Figure 1E). Numerical systematic error was computed as the average absolute value of differences between actual and expected values of responses, measured at, and averaged across each level of expected response. Similarly, numerical imprecision was computed as the average standard deviation of the actual responses measured at and averaged across each level of expected response.

### Entropy and estimation of the number of different responses

We estimated the effective number of different responses used by using the distribution of the provided answer (Figure 6A). While this distribution would provide us with the counts of responses with non-zero probability, some responses may be used very infrequently and affect this measure significantly. To overcome this limitation, we calculated the entropy of the response. More precisely, as the entropy of a uniform discrete random variable with *n* possible outcomes is ln(n), we used the exponential of the entropy of the response as a proxy to measure the effective number of different responses utilized by each pDNN.

### Neural representational similarity (NRS)

We computed the mean response of each neuron for each region individually while receiving each of the distinct 380 operations. Then, we computed the correlation (across neurons) between each of these mean responses, obtaining a 380x380 similarity matrix *M* (shown in Figure 4B) for each region. For practical visualization purposes we sort the rows and columns of *M* so that (1) addition problems come before subtraction problems, (2) among similar type of problems, operations with smaller results come first, and (3) among similar type of problems with the same result, operations with smaller first operands come first. To compute the *NRS* between addition and subtraction problems (referred to as *add-sub similarity*) within a region, we average the similarity between each pair of addition and subtraction problems. To compute the *NRS* between addition problems (referred to as *add-add similarity*), we average the similarity between each pair of two addition problems. To compute the *NRS* between subtraction problems (referred to as *sub-sub similarity*), we average the similarity between each pair of two subtraction problems.

### Geometrical properties of result-manifolds

Recent theoretical advances^58,59^ have defined geometrical metrics that are helpful to understand separable manifolds in neural representations. They quantify the separability of different manifolds using manifold capacity, which measures how easy it is to distinguish two random subgroups of the manifolds. They show that this manifold capacity can be computed from geometrical properties of the manifold, namely the average manifold radius, the average manifold dimensionality, and the correlation between the center of manifolds. Manifold radius reflects the size of the manifold, manifold dimensionality reflects the number of effective dimensions within which the manifold evolves, and correlation between center of manifold reflects the alignment of manifolds. We compute the manifold capacity, the manifold dimensionality, and the correlation between manifold centers for each layer in pDNN separately by using: https://github.com/schung039/neural_manifolds_replicaMFT.

#### Step 7: Comparing effect of hyperexcitability in pDNN vs effect of MLD in children

To compare children and pDNN at the neural level we compared their representational similarity. We focused on representational similarity between addition and subtraction. In prior empirical work ^21^ the correlation coefficients were normalized using the Fisher z-transform before performing group level analysis. Here, to compare pDNN correlation coefficients with the correlation coefficients obtained from fMRI data, we applied the inverse of the z-transform (i.e. the tangent hyperbolic function) on the representational similarity from fMRI data reported in ^21^.

## Acknowledgement

This work was supported by the National Institutes of Health (R01HD059205, R37HD094623), the National Science Foundation under Grant No. (NSF2024856), the Stanford Institute for Human-Centered Artificial Intelligence (HAI), and the Stanford Maternal and Child Health Research Institute (MCHRI).

