## Supplementary Information for "Digital twins for understanding mechanisms of learning disabilities: Personalized deep neural networks reveal impact of neuronal hyperexcitability"

### pDNN model architecture

The pDNN model is based on the CORNet-S architecture (Kubilius et al., 2018). First, V1 receives as input a 28x140 picture with 3 color channels, that is, a 3x28x140 tensor. V1 transforms this input into a 6x7x35 tensor through a chain of a 7x7 convolution with stride 2 augmenting the number of channels to 64, a 3x3 max pooling with stride 2, and a 3x3 convolution, each of the convolution being followed by batch normalization and a ReLU non-linearity. The output of V1 (V2, V3) is then fed as input to V2 (V3, IPS), which transforms it into a 128x4x18 (resp. 256x2x9, 512x1x5) tensor. V2, V3, and IPS are built with the same building block that resembles a recurrent version of a residual block of the ResNet architecture, which has proven to be one of the best performing models on various benchmark datasets in different domains (He, Zhang, Ren, & Sun, 2016). This building block is recurrent, but only the last output it produces is being fed as input to the next block. At time $t = 0$, the input to the block is the last output produced by the previous block, which is first transformed through a 1x1 convolution increasing two times its number of channels. For the following time steps, the input to the block is directly replaced by the feedback of its own output at the previous time step. Moreover, at each time step, the block transforms its input through a chain of 3 convolutions, each followed by batch normalization and a ReLU non-linearity: (1) a 1x1 convolution increasing four times the number of channels, (2) a 3x3 convolution being performed with stride 2 at time $t = 0$, and with stride 1 at all other times, and (3) a 1x1 convolution decreasing four times the number of channels. Furthermore, a skip connection is added between the input before the last ReLU non-linearity which, at time $t = 0$, adds a 1x1 convolution of the input with stride 2 before passing it through the nonlinearity, and at all other times, simply adds the input of the block before passing it through the nonlinearity. V2 and IPS are running for 2 timesteps, whereas V3 is running for 4 timesteps. We focus the analysis of neurons in V2, V3 and IPS on their last timestep. Finally, to produce the output of the whole model, the last output from IPS is fed in a simple linear decoder preceded by an adaptative average pool which enforces the input of this linear decoder to be of dimension 512 (i.e., the number of output channels of IPS). In the original work, the output has 1000 dimensions as there are 1000 different classes in ImageNet. In this work, we only changed the output dimension of that last linear decoder to match the number of classes we consider, that is, 19 dimensions for 19 classes representing each of the 19 different results from 0 to 18 we consider.

**Figure SI 1.Hyper-excitability diminishes differentiations of neural representations across iterations and across pDNN layers.** Evolution of DNN NRS for different levels of neural excitability measured as neural gain *G*. Neural gain *G* is represented by color, varying from blue (*G* = 1) to yellow (*G* = 5). **A-C.** As neural excitability increases, **A.** NRS between addition and subtraction problems (add-sub NRS), **B.** NRS between addition problems (add-add NRS), and **C.** NRS between subtraction problems (sub-sub NRS) show slower decreases with training.

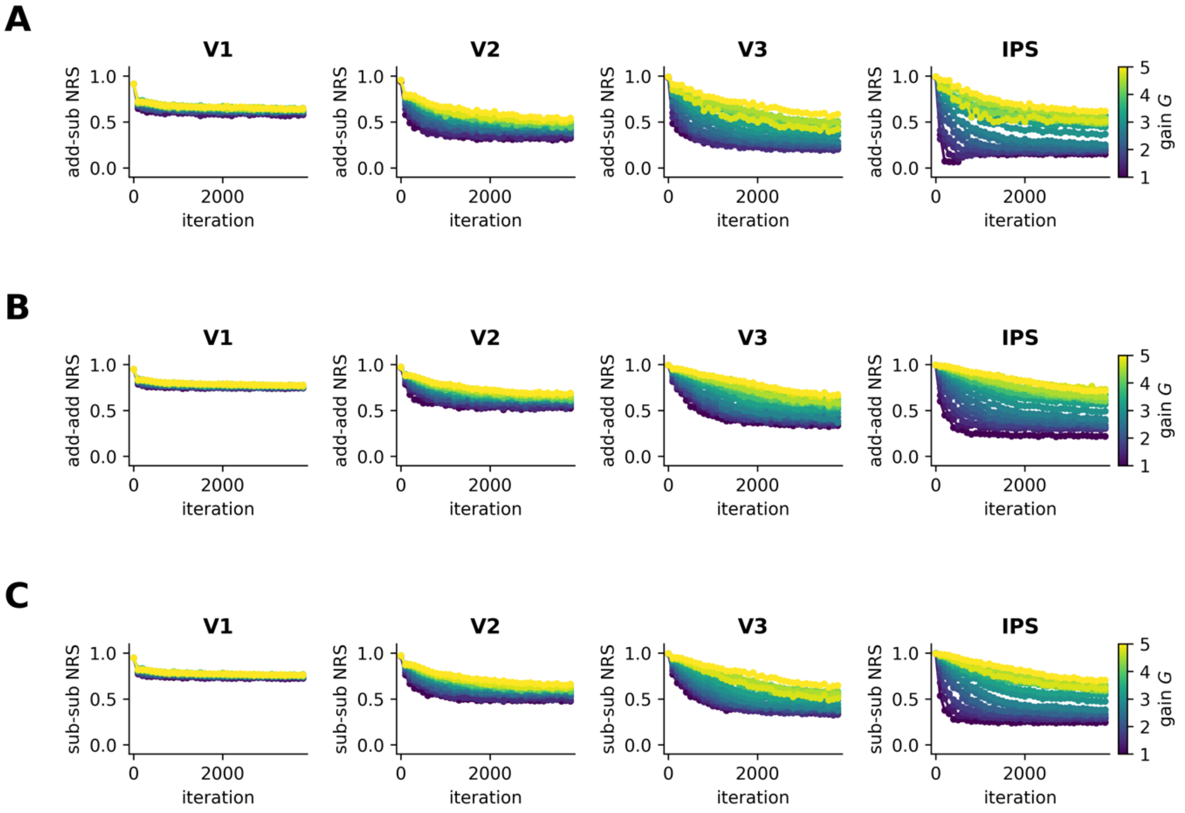

**Figure SI 2. pDNNs predict deficit in neural differentiation between and within operation.** Comparison of NRS across pDNN layers (V1 to IPS) for models matched to children with MLD (red) and typically developing (TD) children (blue). MLD pDNNs show significantly higher **A.** add-sub NRS, **B.** add-add NRS, and **C.** sub-sub NRS compared to TD pDNNs across all layers, indicating both reduced neural differentiation between problem types and within problem types. The effect size of the difference in NRS between MLD and TD pDNNs increases along the network hierarchy, suggesting a more pronounced deficit in higher-order processing regions.

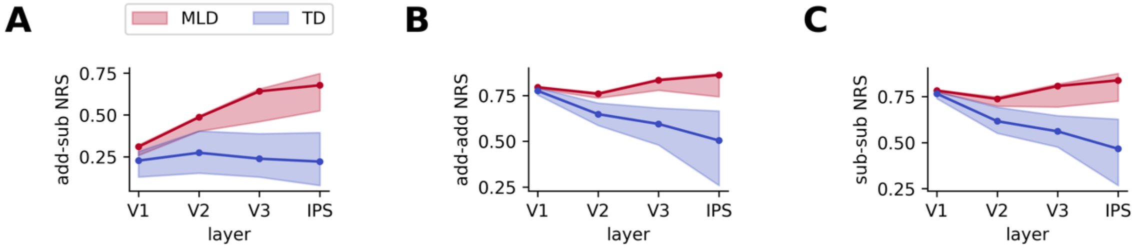

**Figure SI 3. Early deficits in systematic numerical error and numerical precision are overcome with more training.** Evolution of DNN numerical properties for different levels of neural excitability measured as neural gain $G$. Neural gain $G$ is represented by color, varying from blue ($G = 1$) to yellow ($G = 5$). **A-C.** Early in training, as neural excitability increases, **A.** systematic numerical error and **B.** numerical imprecision increase, and **C.** the number of different responses used decrease. All these deficits are overcome with more training, i.e. **A.** systematic numerical trueness, **B.** numerical imprecision, and **C.** number of different responses used reach similar higher levels with additional training regardless of neural excitability.

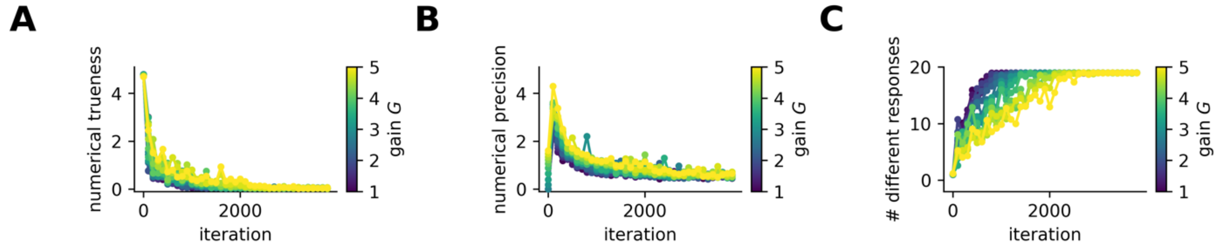

**Figure SI 4. Degradation of manifold geometrical properties of neural representations caused by hyper-excitability are partially overcome with more training.** Evolution of manifold geometric properties for different levels of neural excitability measured as neural gain *G*. Neural gain *G* is represented by color, varying from blue ($G = 1$) to yellow ($G = 5$). **A-C.** As neural excitability increases, **A.** manifold capacity decreases, **B.** manifold dimensionality increases **C.** and correlation between center of manifolds increases, indicating harder to discrimate, more complex and more aligned representations. Not all of these deficits are overcome even with additional training.

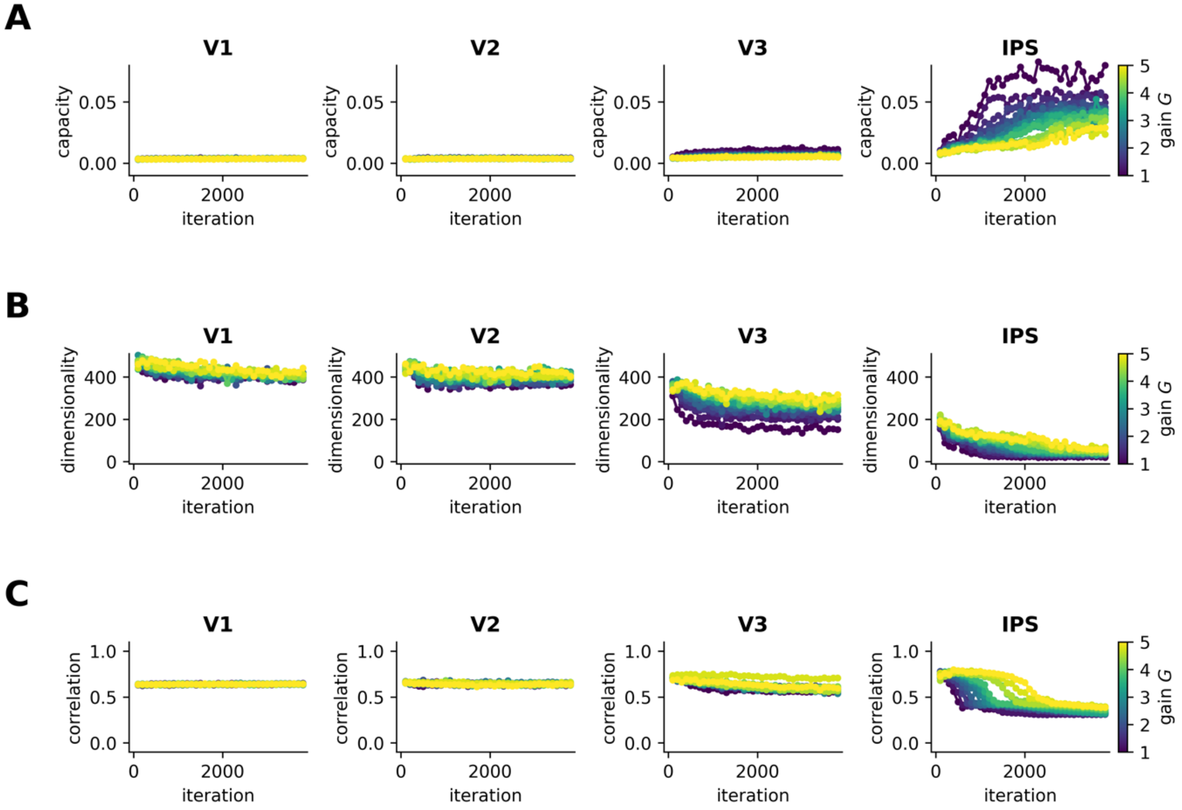

**Figure SI 5. Neural hyper-excitability degrades manifold geometry of latent representations across layers in pDNNs. A-C.** Three key manifold properties in the IPS layer of pDNNs change with neural gain levels across layers. **A.** Manifold capacity, reflecting the separability of neural representations, shows decrease with higher excitation and increase along hierarchy, indicating that hyper-excitability makes it more difficult to distinguish between different numerical manifolds and that it is easier to distinguish between different numerical manifolds higher in the hierarchy. **B.** Manifold dimensionality, indicating the complexity of the representational space, increases with greater neural gain and decreases along hierarchy, suggesting that hyper-excitability leads to more complex and less efficiently organized representations, and that representation are simplified along the hierarchy. **C.** Correlations between manifold centers, relating to the alignment of representations, increase with neural gain and either decrease along hierarchy for smaller gain or increase along hierarchy for higher gain, implying that hyper-excitability causes the centers of different numerical manifolds to become more aligned, potentially leading to increased interference between representations, and that this alignement is amplified along hierarchy for higher gains but reduced along the hierarchy for smaller gains.

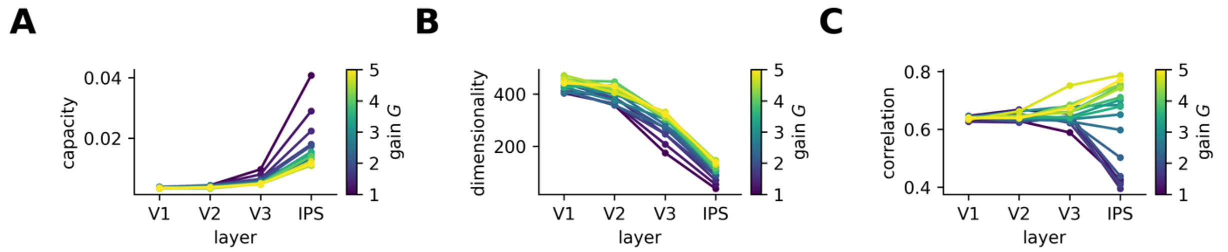

**Table SI 1. Cognitive profiles of participants.** Refer to (Chen et al., Neuropsychologie 2021) Table 1 for a more complete description of the cognitive profiles.

| | **TD children (12F/12M)** | **MLD children (13F/8M)** |
|---|---|---|
| **Age** | 8.40±0.64 | 8.34±0.65 |
| **Full-scale IQ (WASI)** | 111.83±10.26 | 108.33±10.69 |
| **Reading Comprehension (WIAT II)** | 109.04±10.19 | 104.10±9.45 |
| **Numerical Operation (WIAT II)** | 110.75±9.60 | 85.00±4.29 |

**Figure SI 6. Exemples of stimuli of addition and subtraction.**

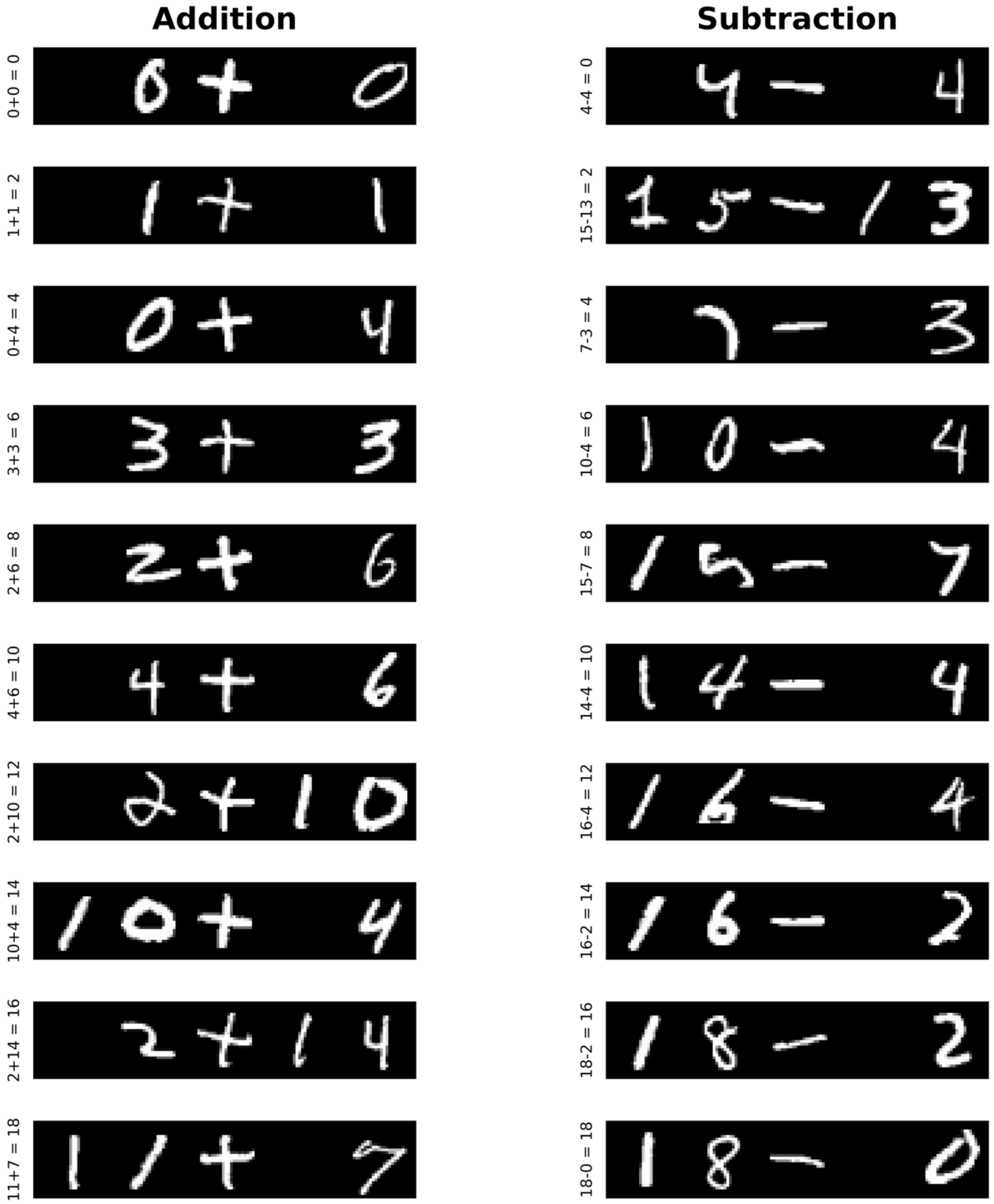
